# Hippocampal-cingulate dynamics in the human brain link reinforcement-learning and memory

**DOI:** 10.64898/2026.09.10.750374

**Authors:** Salman E. Qasim, Fedor Panov, Lizbeth Nunez, Ariane E. Rhone, Hiroto Kawasaki, Christopher Kovach, Christopher Garcia, Brian Dlouhy, Xiaosi Gu, Ignacio Saez

**Affiliations:** Department of Neurosurgery, Rutgers Robert Wood Johnson Medical School, New Brunswick, NJ 08901; Department of Psychiatry, Icahn School of Medicine at Mount Sinai, New York, NY 10029; Department of Neurosurgery, Icahn School of Medicine at Mount Sinai, New York, NY 10029; Department of Neuroscience, Icahn School of Medicine at Mount Sinai, New York, NY 10029; Department of Neurosurgery, University of Iowa, Iowa City, IA 52242; Department of Neurology, Icahn School of Medicine at Mount Sinai, New York, NY 10029; Department of Psychiatry, Yale School of Medicine, New Haven, CT 06511; Department of Biomedical Informatics and Data Science, Yale School of Medicine, New Haven, CT 06511

## Abstract

Reinforcement learning (RL) models describe how reward computations shape our choices, but whether the same computations also shape memory in the human brain is unclear. To address this question, we combined multi-areal intracranial recordings with computational modeling of reward and memory in neurosurgical patients. Patients played a gambling task in which decisions yielded monetary rewards tied to trial-unique images, followed immediately by a recognition test for those images. Model-derived positive reward prediction errors (RPEs) predicted later memory for each image. During feedback, multivariate high-frequency activity revealed the spatiotemporal evolution of RPE representations across distributed prefrontal cortex. During subsequent recognition, however, only anterior cingulate cortex (ACC) transiently reinstated RPE representations, and successful recognition specifically associated with hippocampal reinstatement of these ACC representations. Hippocampal-cingulate theta synchrony scaled with RPE during recognition, alongside hippocampal theta decoding of upcoming memory choices. Thus, the RL computations that guide decision-making also shape memory through hippocampal-cingulate circuit dynamics in the human brain.

## Introduction

Unexpected outcomes are informative: they teach us which actions led to good surprises, and which to bad surprises. These prediction errors—formalized by reinforcement learning (RL) frameworks—matter for both biological brains^1^ and machines^2^. Midbrain dopaminergic activity is canonically correlated with prediction errors^3^, providing a biologically plausible signal for learning adaptive behavior. While this mechanism allows learning of action-outcome associations, less is known about how the actual events surrounding those outcomes are retained in memory. This question is under-explored, partially because RL and memory are often treated as distinct computational problems. In contrast to RL, associated with striatal, prefrontal, and dopaminergic mechanisms for learning statistical regularities to guide value-based decision-making, memory is often associated with medial temporal lobe mechanisms for preserving individual events in a form that can later support recognition, inference, and flexible behavior^4^. But recent work has begun to suggest these systems are not, in fact, independent: reward prediction errors (RPEs) can enhance later memory for stimuli encountered during RL, even when those stimuli are incidental to the task itself^5–11^, and midbrain dopamine enhances hippocampal plasticity mechanisms that support memory encoding^12–14^. In other words: surprising moments are remembered better, but it is unclear how a signal built to guide choices also determines what persists in memory.

The relationship between these processes has important implications. Impaired reward-processing, a key characteristic in many mental health disorders^15–17^, may have important, but poorly understood effects on memory^18, 19^—for example, a maladaptive inability to prioritize positive, rewarding experiences in memory^11^. Answering how reward-mediated learning affects memory processes likely requires moving beyond canonical midbrain-striatal accounts of RL. Whereas reward-related computations are represented broadly across prefrontal cortex^20–30^, the hippocampus is thought to bind items to their context. Linking the two processes, dopaminergic influences on hippocampal function may play a role in encoding surprising events in memory^4,5,12–14, 31, 32^. These observations suggest an interaction between prefrontal representations of outcome significance and hippocampal mechanisms for mnemonic binding and retrieval. However, probing these interactions directly in humans is challenging due to the lack of spatial and temporal resolution needed to resolve the rapid, circuit-level dynamics linking RL and memory using noninvasive human neuroimaging methods.

Here, we tested the hypothesis that RL selectively modulates mnemonic circuits to prioritize events in memory by combining computational modeling with intracranial recordings in humans performing a decision-making task followed by recognition memory. We asked whether trial-wise RPEs predict later memory, how RPE-related information is represented across human prefrontal cortex during learning, and whether these representations reemerge during retrieval in concert with hippocampal dynamics. By tracing reinforcement-learning computations from feedback to later remembering, we sought to identify how a core algorithm of adaptive behavior also helps determine which moments persist in memory.

## Results

### Positive RPEs enhance memory in healthy cohorts and neurosurgery patients with epilepsy

We used a behavioral task to assess the influence that RPEs experienced during decision-making have on subsequent memory (see *Methods*)^11^. This task consists of a two-arm bandit reversal learning game with probabilistic rewards, in which participants are instructed to maximize reward by drawing cards, and each card is associated with a unique image stimulus drawn from a database with normed perceptual memorability ratings (PM). This decision-making task is followed immediately by a recognition memory task utilizing the same stimuli seen during the outcome period of the decision-making task along with novel lure images (Fig. 1A). The task was performed by neurosurgical patients with intracranial electrodes in the epilepsy monitoring unit (*n* = 23, age=37.1 ± 14.9, male=9; see *Methods*) as well as an online cohort of healthy participants (*n* = 208, age = 40.5 ± 12.9, male=104; see *Methods*). Participants from both cohorts were excluded from relevant analyses if performing at or below chance level in the 2-arm bandit task or recognition memory task. Neurosurgical patients learned to pick the most rewarding deck in each block (Fig. 1B, top), suggesting that patients were able to learn the appropriate reward probabilities during each block of the decision-making task. While patients consistently exhibited slower reaction times than the online cohort during decision-making (Fig. 1B, bottom, *t*_217_ = 2.91, *p* = 0.008), they exhibited a similar percentage of optimal choices (Fig. 1B, bottom, *t*_217_ = −1.05), suggesting intact underlying reward computations in the patient cohort. We tested six computational models of behavior in the decision-making task (Fig. 1C, S1, Supp. Table 1; see *Methods*). Specifically, we sought to determine whether this task elicited choices consistent with an RL framework as opposed to heuristic strategies or Bayesian state inference. As in prior work^11^, the Rescorla-Wagner RL model performed the best at modeling participants’ behavior in this task (see *Methods*), suggesting that participants actively track reward outcomes and compare them to expected rewards (Fig. 1D). We validated this behavioral finding and model-fitting in the large online cohort of healthy participants (Fig. S2A-C). We thus used this standard Rescorla-Wagner model to estimate subject-level learning rate and inverse temperature (Fig. 1E), as well as trial-level RPEs (see *Methods*).

**Figure 1:**
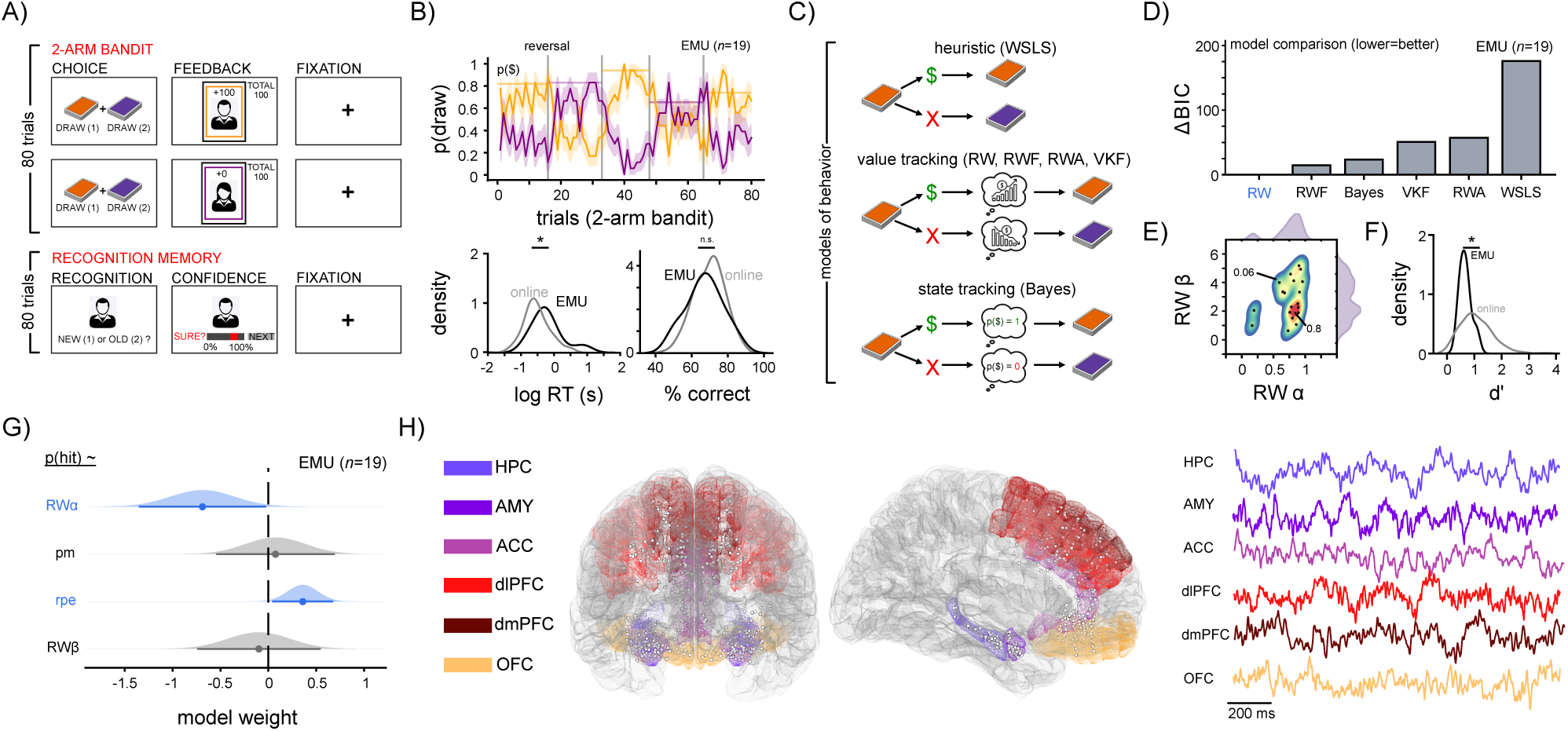
Model-estimated RPEs drive decision-making and predict subsequent recognition memory in neurosurgical patients. A) Top: schematic of two representative trials of the two-arm bandit task. Participants were cued to select a card from one of two decks (orange or purple) on each trial, then provided with feedback and a trial-unique face image, followed by a fixation cross. Bottom: schematic of one trial of the recognition memory task. After having performed all 80 trials of the two-arm bandit task, subjects were cued with face stimuli (old and lure images) and asked to provide a recognition response and confidence rating, followed by a fixation cross. B) Top: probability of drawing from the male (orange) or female (purple) decks as a function of trial and reward block (mean reversal trial indicated by vertical lines). Horizontal lines denote probability of reward for the more rewarding deck on each block. Shaded lines denote a 95% confidence interval. Bottom: Density of log reaction time and percent correct choices for both online (gray) and neurosurgical (black) cohorts. Asterisk denotes a significant difference. C) Schematic of different computational modeling approaches for behavior. D) Model performance, specified by the integrated BIC across six candidate reinforcement-learning models fit hierarchically to subjects’ choice sequences on the 2-arm bandit task. Bar height represents ΔBIC, each model’s integrated BIC relative to the best-fitting model in the set (best = 0); lower is better. E) Joint and marginal distributions of parameter estimates from the RW model, depicting the learning rate (α) and inverse temperature (β). Warm colors indicate higher density. F) Density of memory performance measured by d’ for both online (gray) and neurosurgical (black) cohorts. Asterisk denotes a significant difference. G) Sampling distributions for the fixed effects in a binomial mixed-effects model examining how trial-level reward prediction error (rpe), image memorability (pm), and subject-level Rescorla–Wagner parameters (learning rate α, inverse temperature β) influence recognition memory success. Each curve is a normal approximation to the coefficient’s sampling distribution, and the horizontal bar spans the 95% Wald confidence interval. The dashed vertical line indicates a coefficient of 0. Distributions whose 95% CI excludes 0 are shaded blue, indicating a statistically reliable effect, while those whose CI includes 0 are shaded gray (trial-wise RPE *p* = 0.023; subject-level α *p* = 0.04). H) Left: electrode locations across all 19 participants (white circles), transformed to MNI coordinates, plotted on a schematic glass brain. Functional regions are denoted by color. Right: example LFP traces recorded from each region.

Next, we looked at performance during the recognition test, which occurred immediately after the 2-arm bandit task concluded. Unlike in the 2-arm bandit task, neurosurgical patients exhibited significantly worse recognition memory performance (Fig. 1F, *t*_210_ = −6.1, *p* < 0.001). We next measured the contribution of model-estimated RPEs to subsequent memory, controlling for non-reward (e.g. perceptual) contributions to memory through the memorability ratings associated with each image. Using a mixed-effects regression model that also accounted for subject-level traits such as learning rate (α), and inverse temperature (β), we found that positive RPEs were significantly predictive of subsequent recognition success in the neurosurgical cohort, while subject-level learning rate (α) was negatively correlated with recognition (Fig. 1G, *weight_RPE_* = 0.36, *p* = 0.023, *weight*_α_ = −0.69, *p* = 0.04). Despite the difference in overall memory performance between the neurosurgical patients and healthy participants, we validated that the behavioral phenotype of RPE-mediated memory generalized to the large online cohort (Fig. S2D, *weight_RPE_* = 0.12, *p* = 0.01). We next focused our analyses of neural data on regions critical to decision-making and memory, which were well represented in our patient sample (Fig. 1H, Supp. Table 2): the dorsal medial prefrontal cortex (dmPFC), dorsolateral prefrontal cortex (dlPFC), orbitofrontal cortex (OFC), amygdala (AMY), and hippocampus (HPC).

### RPEs are encoded across prefrontal cortex during feedback

Because RPEs were predictive of subsequent recognition, we sought to identify whether a relationship existed between RPEs and neural activity in decision-making and memory regions. To do so, we computed spectral power (see *Methods*) during the feedback period of the decision-making task—a 1.5 second window of time in which participants receive either a reward or no reward, and are shown the image associated with their selection. This time window has previously been shown to capture cortical encoding of reward computations like RPE^30^. Because high-frequency activity (HFA, 70-200 Hz; Fig. 2A) is a proxy of local ensemble spiking^33^, we hypothesized it would be capable of capturing high-resolution patterns related to the joint encoding of the image stimuli and their associated RPEs, and chose to focus our analysis on this frequency band.

**Figure 2:**
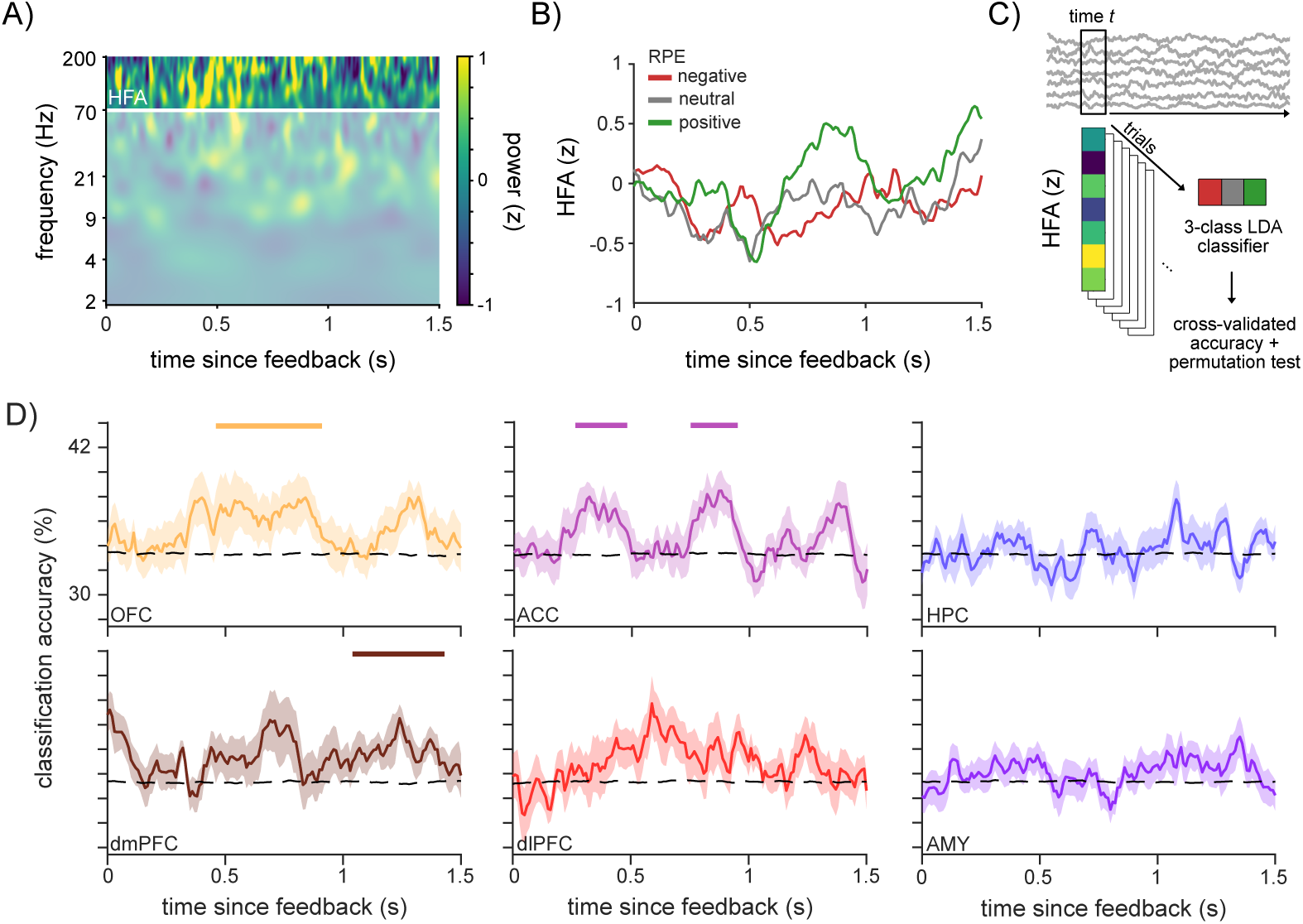
RPE information is encoded across prefrontal cortex, but not medial temporal lobe. A) Time-frequency representation (TFR) of activity in an example OFC electrode during the feedback period of the decision-making task. Warm colors denote increase in power relative to baseline. Trials with negative RPEs (left) show greater theta power (indicated by the red box) compared to trials with positive RPEs (right). B) High-frequency activity in an example anterior cingulate electrode during the feedback period of the decision-making task, split by trial type: negative RPE (red), neutral RPE (gray), and positive RPE (green). C) Schematic of cross-validated multi-class LDA classifier accuracy for RPE class for each region of interest. D) Classification accuracy using HFA as a function of time during reward feedback across all regions of interest. The dotted line denotes the mean of the surrogate values. Solid line denotes mean cross-validated accuracy across subjects. Horizontal lines denote significant time clusters computed using shuffled null data. Shaded lines indicate standard error across subjects.

When participants made choices that resulted in unexpected outcomes during this feedback period, we observed electrodes in prefrontal cortex exhibiting changes in HFA in accordance with these RPEs (example in Fig. 2B). Because electrode locations are not perfectly matched across subjects, and because neighboring electrodes within subjects often showed effects of opposite sign, we pursued a distributed multivariate approach better suited than averaging univariate effects to detect the underlying population code associated with RPEs (see *Methods*). We thus split RPEs into three classes: negative, neutral, and positive (Fig. 2C) and applied cross-validated multivariate classification using HFA^34^, This approach explicitly models the distributed patterns associated with prefrontal RPE encoding—differences in sign, latency, and representational strength^24, 35, 36^—across adjacent populations, allowing us to quantify information represented in the joint activity of electrodes. We found that RPE could be decoded broadly across medial prefrontal cortex regions (OFC, ACC, dmPFC) during the feedback period (Fig. 2D, cluster-based permutation test, *p_OFC_* = 0.022, *ps_ACC_* = 0.027, 0.03, *p_dmPFC_* = 0.02; see *Methods*): early in OFC and ACC, and then dmPFC, at heterogeneous timescales and across spatial distributions (Fig. S3) consistent with observations in nonhuman primates^35, 36^. In contrast, we could not decode RPE from HFA in the memory-related regions: hippocampus and amygdala (Fig. 2D), consistent with work in human neuroimaging^37^. Similarly, we could not decode perceptual memorability (PM) from any of our regions of interest (Fig. S4A); meanwhile, binary reward (i.e. presence or absence of reward) could be decoded from OFC, dmPFC, dlPFC, and amygdala, but not ACC (Fig. S4B, *ps_OFC_* = 0.005, 0.035, *p_dmPFC_* = 0.002, *p_dlPFC_* = 0.005, *p_AMY_* = 0.029). Hippocampal ripples, which have been implicated in model-based reward learning^38^ was not detectably associated with a difference in prevalence between negative, neutral and positive RPEs (Fig. S5). These results show that HFA across multiple prefrontal regions, including OFC, ACC and dmPFC, heterogeneously encode information about RPEs with overlapping but distinct temporal profiles.

### RPEs elicit ACC pattern reinstatement aligned with subsequent recognition

Having identified a broad representation of RPE across medial prefrontal cortex during reward feedback, we next asked if these representations persisted beyond feedback (where they are relevant for decision-making) into the recognition memory period (where they are incidental to the task at hand). To do this, we cross-tested classifiers trained on data from the feedback period on the recognition cue period immediately following the 2-arm bandit task. In this recognition period, half of the same images were presented again amidst novel lures and participants were asked to make recognition choices. We hypothesized that the neural patterns most informative about RPE would be reactivated when RPE-associated images were shown as recognition cues, even in the absence of reward, which would result in significant recognition-period decoding of the RPE-trained classifier. We found that ACC exhibited above-chance cross-decoding during the recognition cue (Fig. 3A; cluster-based permutation test, *p_ACC_* = 0.031), an effect that was consistent across 13/15 participants with ACC coverage (Fig. 3B). The timepoints exhibiting the highest decoding accuracy occurred between 300-640 ms following the recognition cue presentation (Fig. 3A), consistent in time with reports of mid-frontal ERPs associated with familiarity during associative visual recognition^39^.

**Figure 3:**
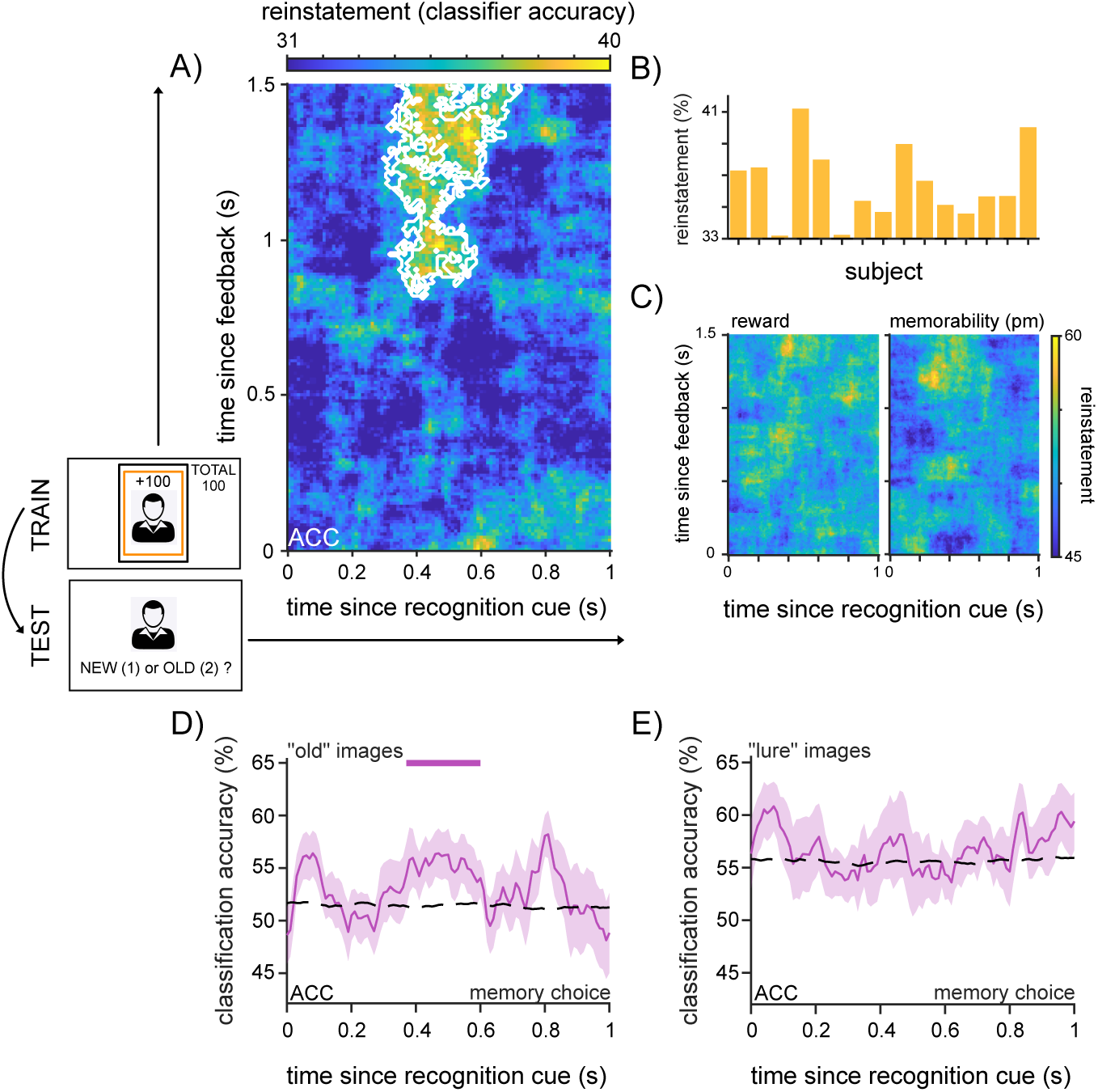
ACC selectively reactivates RPE representations during recognition memory. A) Performance of cross-decoding classifier for RPE class trained on the feedback period (y-axis) and tested on the recognition cue period (x-axis). Warmer colors indicate higher classifier performance. White outline denotes significant time cluster. B) Reactivation in the highlighted cluster for each participant. C) Performance of cross-decoding classifiers trained for binary reward (left) and perceptual memorability (median split, right). Neither classifier yielded significant cross-decoding between the feedback and cue periods. D) Classification accuracy (memory choice during old image cue) using HFA as a function of time during cue period in ACC. The dotted line denotes the mean of the surrogate values. Solid line denotes mean cross-validated accuracy across subjects. Horizontal lines denote significant time clusters computed using shuffled null data. Shaded lines indicate standard error across subjects. E) Classification accuracy (memory choice during lure image cue) using HFA as a function of time during cue period in ACC. The dotted line denotes the mean of the surrogate values. Solid line denotes mean cross-validated accuracy across subjects. Shaded lines indicate standard error across subjects.

In contrast, no other prefrontal region exhibited significant cross-decoding of RPE information during memory (Fig. S6), despite their encoding RPE during the decision task. ACC reactivation was RPE-specific, without significant reinstatement of binary reward outcome or binarized perceptual memorability (Fig. 3C). Similarly, ACC reinstatement of RPE information was not detectably associated with the number of ACC electrodes in each participant, total HFA during feedback in the ACC for each subject, or total HFA during cue in the ACC for each subject (Fig. S7A; spearman correlations=−0.31, 0.06, 0.09, all *ps*> 0.05), or participants’ inverse temperature or learning rate (Fig. S7B; spearman correlations=−0.01, −0.26, all *ps* > 0.05). Next, we assessed whether ACC representations were involved with the recognition memory decisions participants were about to make. We applied the same multivariate classification approach to decode the upcoming memory choice (old vs. new) during the cue period from ACC HFA. Critically, multivariate ACC HFA was able to significantly separate old vs. new memory decisions when the cue image was old (hits/misses), and thus associated with prior RPE information. This occurred in the same time window that ACC representations of RPE information were reactivated (Fig. 3D, *p* = 0.046). In contrast, when novel lures were presented, ACC representations no longer separated upcoming old vs. new memory decisions (false alarms/correct rejections; Fig. 3E). These results suggest that, about 400 ms after a cue is presented, multivariate ACC representations contain information about the RPE previously associated with the cue, and about whether the cue would be successfully recognized or not.

Next, we sought to understand whether the reinstatement of ACC feedback representations was directly linked to memory processes, namely hippocampal-driven retrieval. We hypothesized that if successful memory reflects the recovery of prior internal states, neural patterns elicited at reward feedback should also later reappear during successful retrieval. To examine this question, we used feedback-cue RSA and asked whether reinstatement of representational structure during the memory task distinguished upcoming old vs. new memory choices. We found that within-ACC reinstatement was not detectably associated with a difference between old and new cues (Fig. 4A; cluster-based permutation test, all *ps* > 0.05), despite the fact that ACC reactivated feedback-period RPE representations (Fig. 3A) and ACC activity differentiated old and new choices (Fig. 3D). These results, taken together, argue against a simple account in which reinstated RPE geometry in ACC alone determines recognition choice. To that end, we then assessed feedback-cue RSA in the hippocampus. Within-HPC reinstatement also was not detectably associated with a difference between old and new memory choices (Fig. 4A; cluster-based permutation test, all *ps* > 0.05), arguing against a single-region reactivation account in hippocampus.

**Figure 4:**
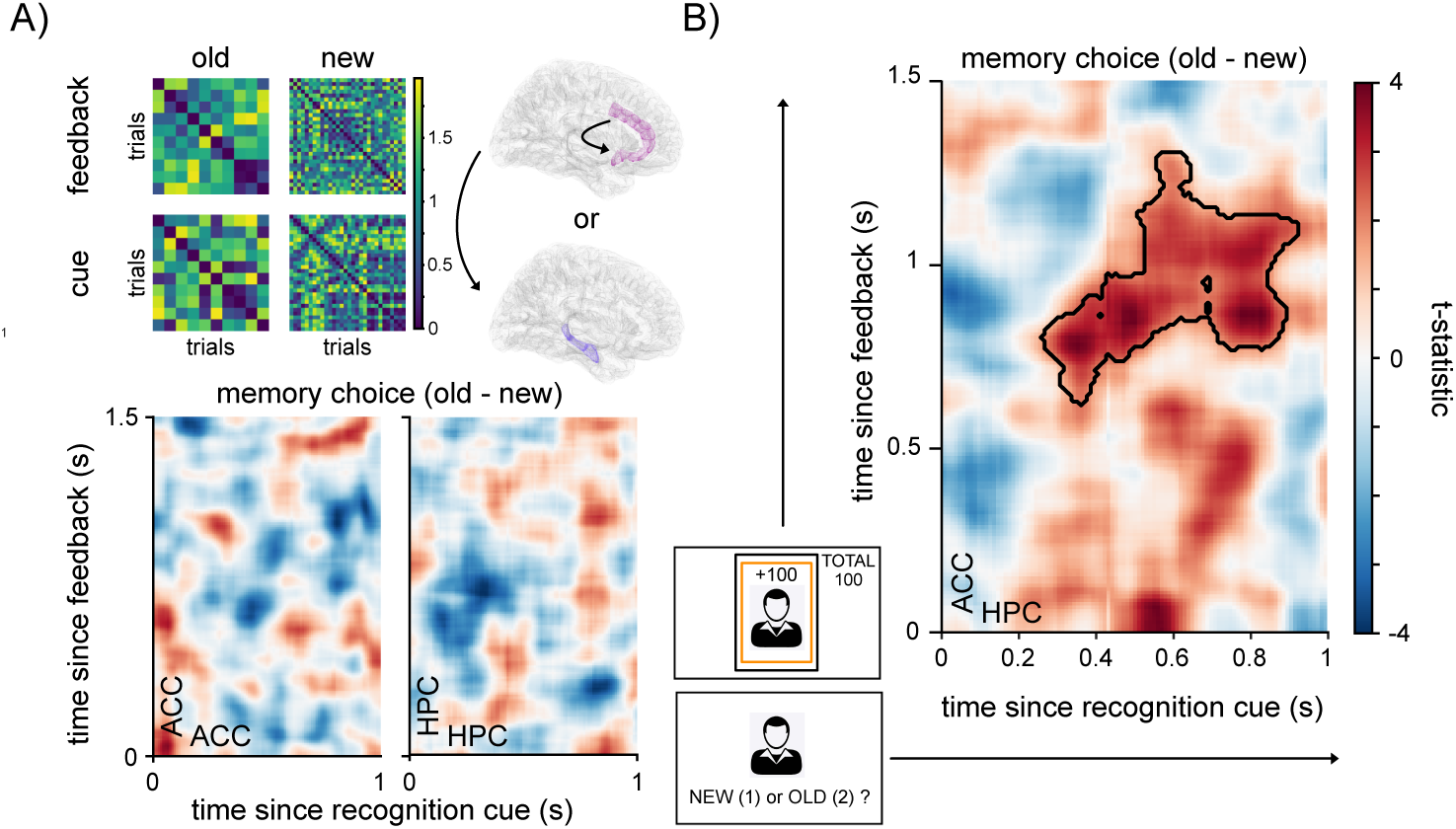
During recognition hippocampus reactivates ACC representations from the feedback period. A) Top: example HFA representational-dissimilarity matrices (RDMs) for cues deemed to be old (left) and new (right) during feedback and cue periods. Schematic denotes two analysis schemes: within-ROI and between-ROI. Bottom: contrast heatmaps of z-scored representational similarity (RSA) for all feedback and cue timepoints (old vs. new) for within-ROI analysis of ACC (left) and HPC (right). Warmer colors denote higher RSA for old choices compared to new choices. B) Contrast heatmap of z-scored representational similarity (RSA) for all feedback and cue timepoints (old vs. new) for between-ROI analysis of ACC feedback to HPC cue. Warmer colors denote higher RSA for old choices compared to new choices. Outline denotes significant cluster using cluster-based permutation test.

### Hippocampal-cingulate coordination integrates RPE representations with memory choice

Because single-region reactivation in either ACC or HPC did not account for recognition memory, we sought to understand whether reinstatement-driven memory choices relied on cross-regional interaction between ACC and HPC. To do this, we performed the same assessment of reinstatement and memory across regions. We found that cross-region reinstatement was associated with successful memory recognition: ACC feedback patterns were more strongly expressed in hippocampal cue-period activity for upcoming old compared to new memory choices (Fig. 4B; cluster-based permutation test, *p* = 0.003). This pattern was exclusive to ACC-HPC reinstatement; hippocampal cue-period reinstatement of other prefrontal regions was not detectably associated with a difference between old and new memory choices (Fig. S8; cluster-based permutation test, all *ps* > 0.05).

ACC-HPC reinstatement during successful recognition was not detectably associated with the number of electrodes in either region (Fig. S9A; spearman correlation=-0.38, -0.1, *ps* > 0.05), participants’ inverse temperature or learning rate (Fig. S9B; spearman correlation=0.04, -0.05, *ps* > 0.05), or overall performance in either the memory (Fig. S10; spearman correlation=-0.07, *ps* > 0.05). Nor did the level of ACC reinstatement of RPE related patterns predict the ACC-HPC reinstatement during successful recognition (Fig. S11; spearman correlation=-0.35, *p* > 0.05). In summary, successful recognition was not simply determined by local recurrence of prior cortical representations of RPE information, but by their cross-regional re-expression in hippocampal retrieval-state activity.

Next, we examined whether hippocampal and ACC theta dynamics were coordinated during memory recognition, given the proposed role of hippocampal theta in memory; we hypothesized such a role could be involved in binding the images to their associated RPEs^40–42^. We started by examining theta power in ACC and HPC contacts. We found that both ACC and HPC exhibited clearer theta oscillations during the memory recognition cue period than during the feedback period (Fig. 5A, B; mixed-effects linear regression, *p_ACC_*, *p_HPC_* < 0.001). Given the transient nature of human hippocampal theta oscillations^43^, we next computed whether the prevalence of transient theta bursts (see *Methods*) differed between RPE classes to assay RPE-mediated changes in theta power. We found no difference between conditions in either HPC or ACC (t-test, all *ps* > 0.05), suggesting that the prevalence of theta bursts does not reflect RPEs. In contrast, multivariate classification of memory choice using hippocampal theta power revealed that hippocampal theta power could differentiate upcoming old from new choices during the presentation of old cues (Fig. 5C; *p* = 0.007); an effect that was absent in ACC theta. This helped to confirm the unique involvement of hippocampal theta in memory recognition, and suggested further investigation into the specific role of hippocampal theta oscillations in coordinating ACC representations.

**Figure 5:**
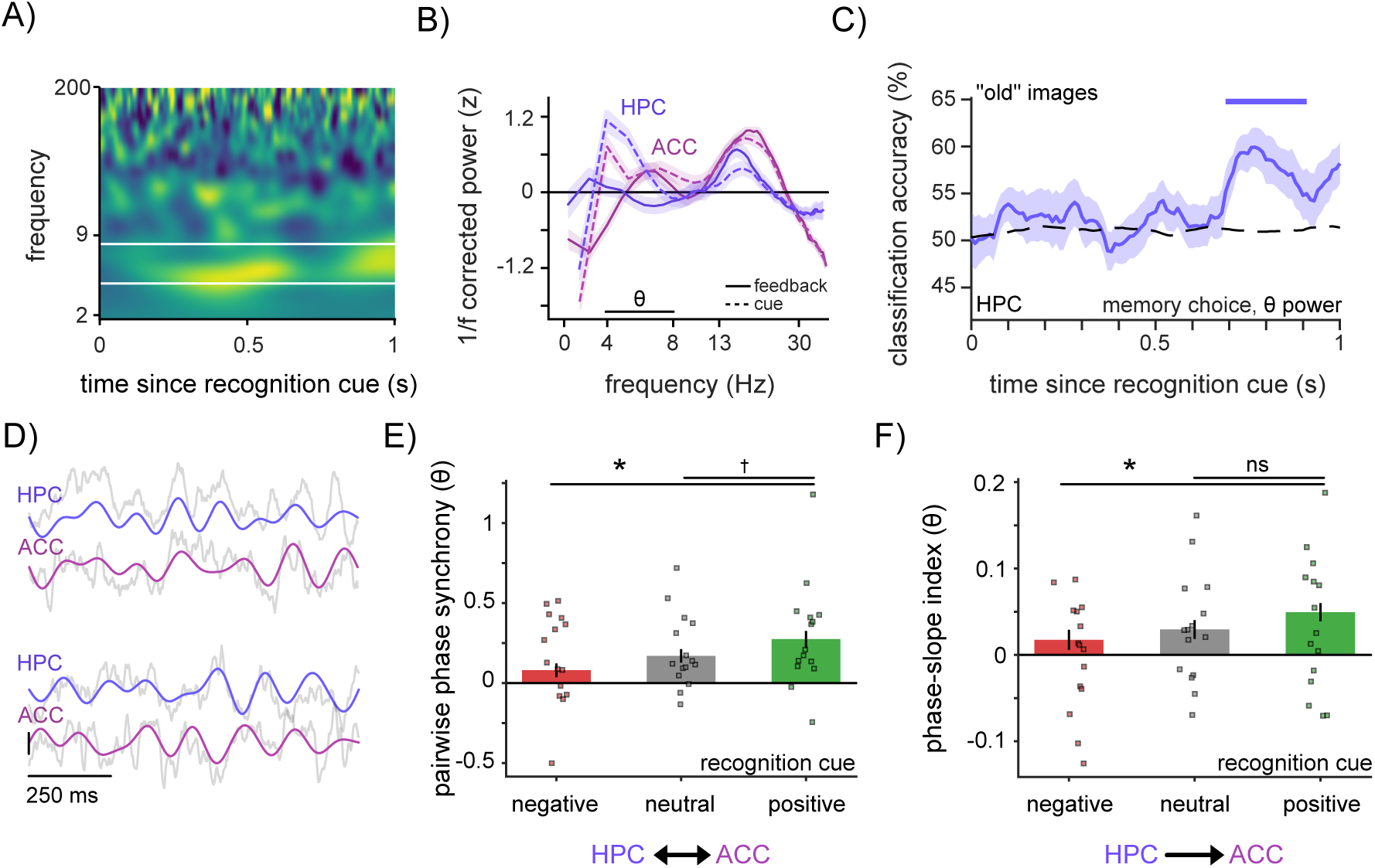
Hippocampal theta dynamics linked to both RPE and memory across hippocampal-ACC circuit. A) Time-frequency representation (TFR) of activity in an example ACC electrode during the cue period of the recognition memory task. Warm colors denote increase in power relative to baseline. B) Z-scored power spectra for all ACC and HPC electrodes following the subtraction of fitted 1/f background spectra, for both the feedback (solid line) and cue (dotted line) periods. Shading denotes standard error across electrodes. C) Classification accuracy (memory choice during old image cue) using theta power as a function of time during cue period in hippocampus. The dotted line denotes the mean of the surrogate values. Solid line denotes mean cross-validated accuracy across subjects. Horizontal lines denote significant time clusters computed using shuffled null data. Shaded lines indicate standard error across subjects. D) Exemplar trials demonstrating hippocampal theta (4-8 Hz) exhibiting a consistent phase relationship with anterior cingulate theta. E) Comparison of z-scored theta pairwise phase consistency (PPC), a measure of undirected phase synchrony, between HPC and ACC during the cue period, for images associated with positive RPEs (green), neutral RPEs (gray), and negative RPEs (red). Bar heights indicate the mean over subjects (dots), with asterisks denoting a significant difference (mixed-effects linear regression, *p_pos_*_:*neg*_ = 0.002, *p_neutral_* = 0.09). F) Comparison of z-scored theta phase-slope index (PSI), a measure of directed phase synchrony, between HPC and ACC during the cue period, for images associated with positive RPEs (green), neutral RPEs (gray), and negative RPEs (red). Bar heights indicate the mean over subjects (dots), with asterisks denoting a significant difference (mixed-effects linear regression, *p_pos_*_:*neg*_ = 0.04, *p_pos_*_:*neutral*_ = 0.2).

Given the role of hippocampal theta phase alignment in associative recognition memory in humans^44^, and theta synchrony between prefrontal cortex and medial temporal lobe in human memory retrieval^45^, we next sought to examine the timing and directionality of HPC/ACC theta activity. We tested whether theta oscillations in the hippocampus reliably synchronized with those in the anterior cingulate during memory recognition (Fig. 5D; see *Methods*). We observed that undirected theta synchrony between HPC and ACC increased for cues previously associated with positive RPEs compared to negative RPEs (Fig. 5E; mixed-effects linear regression, *p_pos_*_:*neg*_ = 0.002, *p_pos_*_:*neutral*_ = 0.09). A directed phase synchrony analysis revealed that HPC theta led ACC theta during positive RPEs compared to negative RPEs (Fig. 5F; mixed-effects linear regression, *p_pos_*_:*neg*_ = 0.04, *p_pos_*_:*neutral*_ = 0.2). Notably, we did not observe significant theta-HFA phase-amplitude coupling within or between HPC and ACC (Fig. S12A-B; mixed-effects linear regression, all *ps* > 0.05), suggesting that this coordination does not arise from a consistent, phase-locked modulation of local ACC high-frequency activity by hippocampal theta, but could instead reflect a more distributed temporally flexible routing of information across the network. Furthermore, the degree of subject-meaned HPC-ACC theta PPC did not predict the magnitude of ACC reactivation of RPE information or ACC-HPC reinstatement during successful recognition—suggesting a degree of independence between the hippocampal-ACC dynamics across slow and fast frequencies as both are engaged in RPE representation and memory-predictive activity (Fig. S13; spearman correlation=-0.33, 0.43, all *ps* > 0.05). Together, these results show that hippocampal-ACC theta synchrony is enhanced for RPE-associated memories and exhibits a directional asymmetry in which hippocampal theta leads ACC theta, but this is statistically independent of the magnitude of neural reinstatement.

## Discussion

How does a signal that guides our choices shape what we remember? Here, we used intracranial recordings to examine the relationship between reward learning and memory directly in the human brain. RPEs, the primary teaching signal of RL, predicted which experiences were later remembered and evoked prefrontal representations that were selectively reinstated by the ACC and the hippocampus. Computational modeling identified RPE-mediated memory in both healthy and neurosurgical cohorts, and direct-brain recordings traced the spatiotemporal evolution of RPEs in the human brain from encoding to retrieval as they help determine which experiences are preferentially retained and later recovered^31, 32, 46, 47^.

These value computations are closely linked to phasic dopaminergic signaling, providing a biologically plausible teaching signal for value updating at the synaptic level^3^. While classically localized to midbrain–striatal circuitry, convergent evidence now indicates that the prefrontal cortex represents many such computations utilized in RL, likely in interaction with corticostriatal loops^48, 49^. Consistent with this view, we were able to demonstrate above-chance decoding of RPEs broadly across prefrontal regions including the OFC, ACC, and dmPFC. The onset time of OFC and ACC RPE-encoding in our task is aligned with the timescale of putative dopaminergic input in non-human primates^50^ and recent intracranial voltammetry work in humans demonstrating dopamine changes in ACC during instrumental learning^51^. One possibility is that prefrontal cortex represents a readout of RPE downstream of midbrain dopaminergic input.

Alternatively, we propose that rather than providing a simple readout of midbrain-striatal dopamine dynamics, the spatial and temporal organization of this decoding is consistent with human neuroimaging studies implicating distributed and diverse frontal value signals^20–23, 52^ and with nonhuman primate recordings showing heterogeneous value and decision coding across orbitofrontal, cingulate, and lateral prefrontal cortex^24–26, 53–55^. Recent human iEEG studies similarly suggest that ACC, dlPFC, dmPFC, and anterior insula carry temporally specific but partially dissociable signals related to signed RPEs, unsigned surprise, expectancy violations, and behavioral salience^28, 29, 56^. Further evidence from OFC showed that local high-frequency activity simultaneously tracks expected value, experienced outcome, and updating-related variables, consistent with OFC maintaining a rich task-state model rather than merely signaling reward magnitude^28, 30^. The temporal resolution of iEEG further characterizes these distributed signals not only anatomically but also by their precise temporal organization across task events. In our data, for example, specific patterns of ACC HFA associated with encoding were subsequently reactivated at specific times during retrieval, revealing structured temporal relationships between neural representations across distinct stages of behavior. Such temporal structure provides constraints on the sequence and timing of computations within the underlying circuit, and therefore on when and where neuromodulatory influences could potentially shape those computations. Thus, multivariate RPE decoding may reflect more than a simple RPE readout redundant with the midbrain-striatal circuit; instead, the spatial and temporal diversity we observe may reveal distributed circuit dynamics through which prediction-error information is incorporated into, and later re-engaged by, memory-guided behavior.

Could one of those computations be related to the memory enhancement associated with these RL computations? Most research linking RL and memory focuses on how memory informs learning and decision-making processes^57, 58^, and the benefits such integration would confer on RL^59^. But recent work has begun to demonstrate how RL confers memory benefits for stimuli associated with the RPEs^5–11^. The RPE-mediated memory in these instances, and that we observed, is separate from RL—for example, individuals would plausibly show long-term learning for the choices that yielded the most positive RPEs. It is also separate from associative learning—although participants learned changing reward contingencies associated with face category, individual faces were encountered only once. Thus, RPEs generated during RL could not act by strengthening associative learning mechanisms for individual face identities; rather, they could modulate the encoding of the episodic representations formed for those one-shot events. Ultimately, the specific context associated with the feedback itself (in this case, an image stimulus) has no utility and is in fact totally irrelevant to the 2-arm bandit reversal learning task. Furthermore, the RPE-mediated memory we observed is likely not due to arousal/affect mediated memory^32^. Why would reinforcement-learning influence memory for incidental stimuli associated with reward computations?

One possibility is that the prefrontal cortex integrates prediction errors with surrounding contextual features during encoding^60^. Storing these context-bound outcome traces could support model-based inference beyond simple model-free action-value updates^40, 60^. Alternatively, reinstating these context-coupled representations could facilitate credit assignment across delays or intervening events, helping connect unexpected outcomes to their specific underlying choices^61^. This possibility would require both the formation of RPE-related representations that persist beyond the immediate learning event and interactions with the hippocampus, a general integrator of stimuli and context^4^. The first requirement aligns with longstanding views of ACC as uniquely positioned to represent reward history over prolonged timescales, even across trials^25, 62, 63^. Across lesions, electrophysiology, and behavior, the ACC consistently emerges as a structure that converts reward prediction errors into adaptive control over ongoing behavior when choices must be sustained, sequenced, or revised over time^35, 50, 64^, in contrast to OFC’s canonical role tied to immediate stimulus/outcome valuation^25, 65, 66^. In fact, recent work has demonstrated that during a three-arm bandit task ACC ensembles exhibited both reward coding and representational drift (a potential biomarker of temporal information and mnemonic function) arguing in favor of a role for contextual representation in ACC reward computations^67^. Indeed, our results highlight that only ACC reinstates RPE-mediated neural patterns during retrieval, in favor of the hypothesis that ACC is unique among prefrontal regions in tracking reward computations on a longer timescale. Critically, hippocampal-ACC theta synchrony was enhanced when memories associated with positive RPEs were cued for recognition.

However, we found that ACC reinstatement is not, itself, a predictor of memory, consistent with the observation that cingulotomy patients did not exhibit memory deficits^68^. Instead, the hippocampus is often positioned as the binding system that links item information with spatial, temporal, and task context into conjunctive memories that can later support recognition, recollection, and flexible inference^69^. During retrieval, the hippocampus is thought to reinstate distributed cortical patterns (such as those in ACC) present during the original event via pattern completion, allowing partial cues to recover broader event representations stored across neocortex^70, 71^. In humans, fMRI evidence likewise suggests that RPEs are associated with increased memory-related hippocampal BOLD activity^5^.

Our findings are therefore supportive of the notion that ACC reinstatement may reflect persistence of a cortical record that an outcome was surprising or behaviorally significant, but with hippocampal recruitment playing a role in reactivating the full episode in which the outcome occurred, including other contextual, potentially relevant features. Our finding that hippocampal reinstatement of ACC feedback patterns distinguished successful recognition is therefore consistent with the idea that hippocampus retrieves the contextual structure attached to reinforcement signals and re-expresses those representations when the incidental stimulus is encountered again. Our findings would thus position the hippocampus as a site where prediction errors acquire mnemonic consequences: not by computing reward prediction errors itself, but by selectively embedding surprising outcomes into relational memory networks that can later guide inference, generalization, and future decisions. One limitation of our study is that we were not able to differentiate what the hippocampus was reinstating: item-level information, or RPE context. Our results suggest a mixture of both, and future work with larger trial counts will be able to measure their relative contributions.

Our findings raise an important question for future research: what is the role of consolidation in linking RL and memory? Consolidation has long been thought critical to the interactions between prefrontal cortex and hippocampus for adaptive memory^31, 41, 72–74^. ACC, in particular, is thought to be critical to long-term, consolidation-dependent memory^75^. During such remote recall, the theta synchrony between ACC and hippocampus increases over the course of consolidation, correlated with the dependence of contextual memory on ACC^76^. We hypothesize, then, that at longer time-scales, the ACC’s reactivation of RPE information (and concurrent hippocampal theta synchrony) would be able to distinguish successful memory, while hippocampal reactivation of ACC representations would become more weakly related to successful memory. While many studies utilizing cohorts of neurosurgical patients with epilepsy present challenges to generalization in the general population, we show that the behavioral–computational phenotypes exhibited by the healthy population are preserved in epilepsy patients, who show comparable task performance, RL parameters, and RPE–memory relationships despite documented impairments in decision-making and memory^77, 78^. This suggests that the neural correlates of RPEs, and their modulation by mnemonic processes, may also be conserved in a general population.

This account provides a mechanistic foundation for potential future therapeutic approaches targeting memory disorders and psychiatric symptoms associated with maladaptive memory bias. RL processes are altered in depression and rumination^16, 79, 80^, and depressive state can modulate how reward prediction errors influence memory formation^11, 81^. These findings motivate future investigations into whether hippocampal-cingulate mechanisms that couple RL signals with memory can be leveraged to modulate memory biases relevant to mental health. Consistent with this possibility, prior iEEG work has demonstrated altered reward-related ACC dynamics in depression^82^, and direct stimulation of the subgenual cingulate has shown promise for alleviating depressive symptoms^83^. Our prior work further demonstrated that stimulation can alter arousal-mediated memory processes in the hippocampus and amygdala by modulating associated increases in HFA^84^. Together, these findings motivate future studies testing whether stimulation of hippocampal-cingulate circuits can selectively influence the relationship between RL and memory, with potential applications spanning mental health and memory disorders. More broadly, by identifying mechanisms through which the human brain endogenously strengthens memory, this work may inform future approaches for enhancing memory function in the context of aging and dementia.

In summary, our findings suggest that RL signals do more than update future choices: they also help determine which moments are carried forward in memory. This is achieved by coordinating anterior cingulate representations of RPEs with hippocampal mechanisms for mnemonic binding and retrieval. By tracing these computations from reward feedback to later recognition, we identify the human ACC–hippocampal circuit as a candidate pathway through which endogenous value signals prioritize experience, linking adaptive decision-making to the selective persistence of memory. This novel understanding of how reward, memory, and cingulo-hippocampal circuit dynamics interact may offer new future opportunities to recalibrate maladaptive memory bias in psychiatric illness and to preserve memory function in aging and neurodegeneration.

## Methods

### Data collection and participants

The EMU study was approved by the Institutional Review Boards at the Icahn School of Medicine at Mount Sinai (IRB 22-00529) and the University of Iowa (IRB 201911084). Patients with treatment-resistant epilepsy who underwent neurosurgical mapping of their seizures provided written consent to participate in behavioral research during their hospital stay. We ensured that a) research participation was voluntary at all stages, and b) that research did not interfere with the integrity of participants’ clinical treatment or experience, in accordance with ethical practices for conducting intracranial research with humans^85^. A total of 23 individuals (9 male, age = 37.1 ± 14.9) participated in the study, and we excluded three participants whose memory performance was at or near chance (*d*^′^ ≤ 0.25), and one participant for technical issues during data acquisition, resulting in a final cohort of 19 participants.

The online study was approved by the Institutional Review Board at the Icahn School of Medicine at Mount Sinai. Participants were recruited from Prolific (http://prolific.co), an online survey platform. A total of 208 individuals (male=104, age = 40.5 ± 12.9) participated in the study. We excluded 8 participants whose overall behavioral performance in the decision-making task was at or near chance from all analyses. We subsequently excluded 7 additional participants whose memory performance was at or near chance from recognition memory analyses. Participants were paid a fixed rate with a bonus computed as a function of reward accumulated in the first task in the experiment. Target sample size was determined based on the results of similar studies investigating the effect of RPE on memory^7,8^. The study was not pre-registered.

### Task

Participants performed an experiment with two distinct stages: a decision-making task, followed by a recognition memory task. The decision-making task was a two-arm bandit task in which participants attempt to maximize their rewards by learning on one of two possible options (decks of cards) on each trial, for a total of 80 trials. Each option was associated with a specific win probability (either 0.8 or 0.2) which reversed four times, every 16 ± 1 trials. As a result, each option’s win probability was always negatively correlated. However, in contrast to traditional bandit/reversal tasks, participants were shown a unique image stimulus after making each choice, associating each reward outcome with a specific image stimulus. These image stimuli were memorable faces drawn from a database on perceptual memorability, where each image was associated with a normed d’ score^86^ that measured how well these images are recognized in a large population sample^87, 88^. Upon completing the decision-making task, participants immediately began a recognition memory task in which 40 of these image stimuli were randomly selected as recognition cues, in addition to 40 novel lure images drawn from the same memorability database with matched d’ scores. The remaining 40 images encoded during the decision-making task were used as recognition cues 24 hours later—these data are not included as a part of the current study. During the recognition memory task, participants were instructed to indicate whether the image was “old” or “new”, and then asked to indicate their confidence in their selection.

### Computational modeling

We used hierarchical modeling to fit six candidate cognitive models to participant behavior in the decision-making task. Subject-level parameters were estimated with hierarchical expectation-maximization under a Laplace approximation (pyEM)^89, 90^: subject-level maximum a posteriori estimates are obtained given a group-level prior, and the group mean and covariance are then updated from those subject posteriors (empirical Bayes). This partial pooling of data across subjects gives more stable parameter estimates and better predictive performance than fitting subjects independently, while still supporting inference on group-level effects^91–96^.

We first fit all six models to our cohort of 19 epilepsy patients tested in the epilepsy monitoring unit (EMU), testing different theories about the mental computations underlying task performance. Four of the models shared a common value-caching architecture, in which value is updated on every trial by a reward-prediction error and choices follow a softmax over those cached values: a Rescorla-Wagner model (RW) with a fixed learning rate; a Rescorla-Wagner model with a fictive learning component (RWF); a valence-asymmetric Rescorla-Wagner model (RWA) with separate learning rates for better- and worse-than-expected outcomes; and a volatility Kalman filter (VKF), in which the learning rate is not fixed but instead adapts, trial by trial, to an online estimate of environmental volatility^97^. To address the possibility that participants were instead using strategies that do not require caching and updating a value for each choice, we also fit a Bayesian change-point filter model that tracks the posterior over which deck is currently better; (*p_switch_*) is the fitted hazard rate, with reward probabilities fixed to the task’s true contingencies^98^; and a win-stay/lose-shift model (WSLS) with a noise parameter governing the probability of repeating or switching choices following wins and losses. These models and their parameters are detailed in Table 1.

**Table 1:** Models, along with free parameters, are utilized for model comparison. The bold row denotes the winning model.

| **model** | **parameters** |
| --- | --- |
| WSLS | $\epsilon$ |
| **RW** | $\alpha, \beta$ |
| RWF | $\alpha, \beta$ |
| RWA | $d, \beta$ |
| VKF | $\omega, \beta$ |
| Bayes | $p_{\text{switch}}, \beta$ |

**Table 2:**
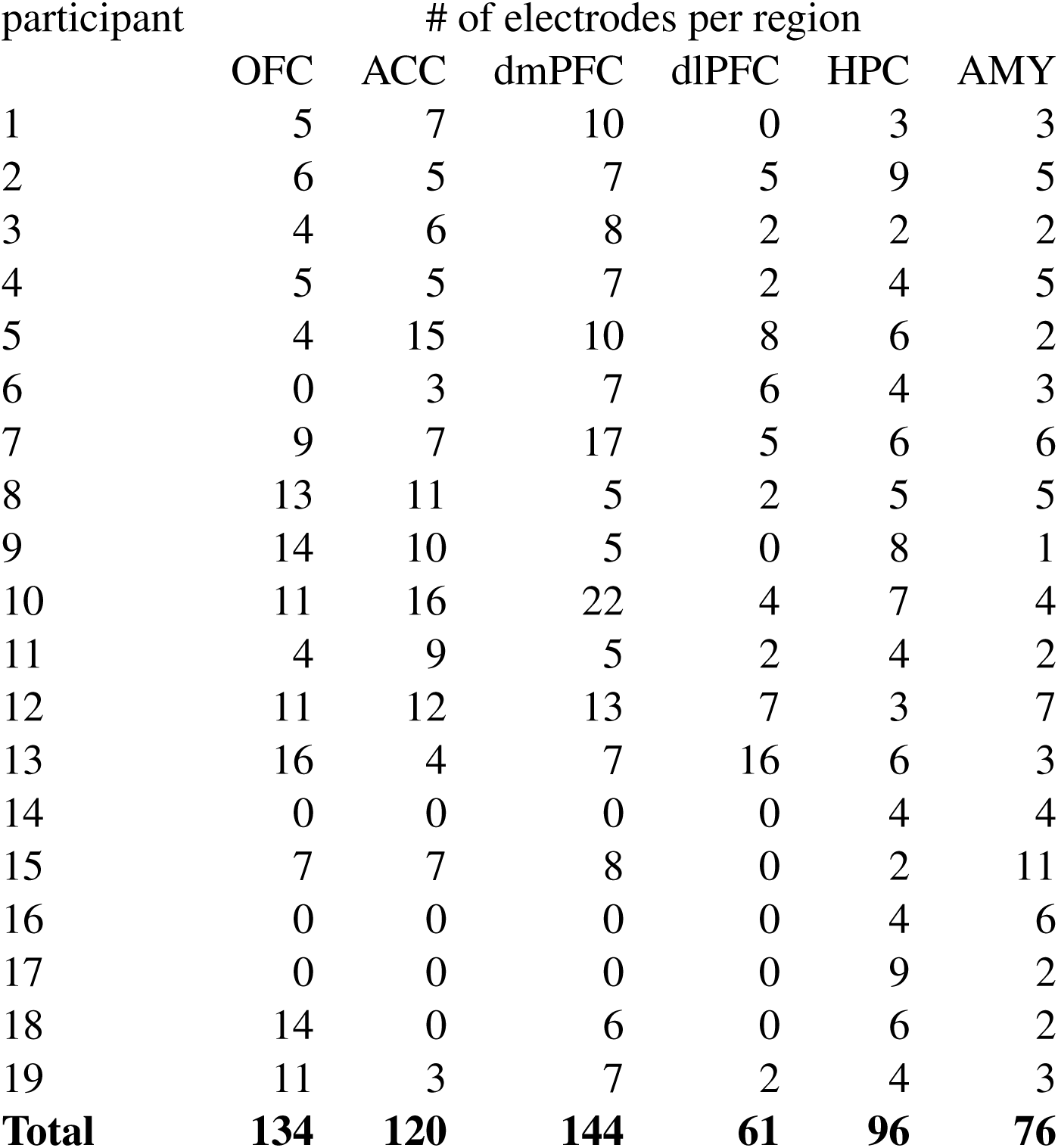
Participants and number of electrodes per region.

#### Model comparison

Computational models were compared using pyEM’s integrated Bayesian Information Criterion (BIC*_int_*; lower values indicate better fit)^89^. The model which performed best was the Rescorla-Wagner RL model (RW). In the Rescorla-Wagner model, reward-prediction errors (RPEs) modulate a learning rate (α) parameter, and RPE-based decisions are determined by an inverse-temperature parameter (β) modulating a softmax choice function. The learning and decision rules for this model are described by the following equations^99^:

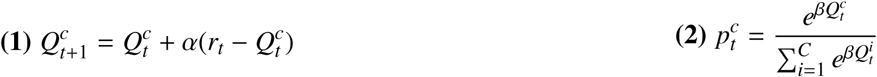

where 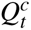 is the value of the chosen option on trial *t*; *r_t_* is the reward received on that trial, and 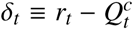 is the model-estimated reward-prediction error (RPE) that determines 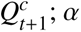; α indexes how quickly values are updated, and β indexes how deterministically those values drive choice.

The remaining five models modify this update or replace it with a different rule entirely. RWF also updates the unchosen option, by the opposite of the experienced prediction error,

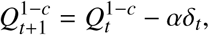

so the value gap between options moves roughly twice as fast as under standard RW for a given α. RWA instead updates the chosen option with one of two learning rates depending on the sign of the prediction error,

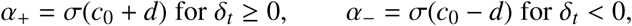

where *c*_0_ = logit(α̅) and the baseline rate is fixed at α̅ = 0.35, so *d* captures any asymmetry in learning from better- vs. worse-than-expected outcomes. The Bayesian change-point filter tracks a posterior belief over which deck is currently more valuable and updates it with a Markov switch hazard *p_switch_*, choosing via a logistic function of that belief—an ideal observer that knows the task’s reward probabilities but has to infer when reversals occur. The VKF keeps the same value-caching architecture as RWF, updating both arms on every trial, but scales each trial’s prediction error by a Kalman gain that reflects the model’s own uncertainty about the value, rather than a fixed learning rate. A free parameter, ω, sets the observation-noise term that shapes this gain and the model’s initial uncertainty; the rate at which the underlying volatility estimate itself evolves is fixed rather than free in this reduced variant, so choices become more or less sensitive to recent outcomes as that volatility process unfolds. WSLS, finally, requires no value representation at all: it repeats a rewarded choice with probability 1 − ɛ/2 and switches away from an unrewarded one with the same probability, with ɛ capturing residual choice noise.

#### Model fitting

Free parameters were fit on an unconstrained latent (Gaussian) scale and mapped to their natural scale via link functions: a logistic map for parameters bounded in (0, 1) (α, ɛ, *p_switch_*), a scaled logistic map for β ∈ (0, 20), and a softplus map for the strictly positive VKF terms; the RWA asymmetry *d*, being unbounded, required no transformation and was fit on the latent scale directly. We initialized group-level means and variances at task-plausible values rather than an uninformative prior, to stabilize the first E-step. At each EM iteration, subject-level maximum a posteriori estimates were obtained given the current group-level prior, and the group-level means and variances were then updated from those subject posteriors; we iterated until the relative change in summed negative posterior log-likelihood across subjects fell below 1 × 10^−5^.

#### Parameter recovery

For each model, we simulated 100 agents using the same trial schedule as the task, drawing parameters from task-plausible hierarchical populations matching the initialization priors. To guard against the EM loop converging to a local optimum, each model was fit from three independent seeds, and the fit with the lowest negative posterior likelihood was retained. We found strong Pearson’s *r* correlations between the true and recovered parameter values for every model (all *r* > 0.5), with the hierarchical refit converging for every model, suggesting our task and models were well suited to estimate these parameters. Because the RW model performed best in our cohort of 19 epilepsy patients, and its parameters were recovered successfully, we next asked whether the same result held in a much larger, independent sample. We re-fit all six models, using the identical pipeline and priors, in 200 healthy online volunteers recruited through Prolific. RW again outperformed the other five models by BIC*_int_* in this online cohort, replicating the result from the EMU sample.

### Data acquisition

Epilepsy patients were stereotactically implanted with depth electrodes (Adtech, Medical, Racine, WI) whose number and location were exclusively determined by the clinical team based on each patient’s clinical needs. Electrophysiological data were recorded using either a clinical monitoring system (Natus, Middleton WI) or an Atlas research recording system (Neuralynx Inc., Bozeman MT) at a sampling rate of 1,000 Hz. At the time of acquisition, LFPs were either referenced to a skull screw or to a 4-contact subdural strip electrode.

### Neural pre-processing

After acquisition, LFP data were bipolar re-referenced, notch-filtered for 60 Hz line noise and harmonics, and re-sampled to 500 Hz for storage. To systematically identify interictal discharges (IEDs), we a) band-pass filtered the LFP between 25-80 Hz, b) rectified the amplitude of the filtered data, c) z-scored the resulting signal, and d) identified peaks in this signal that were at least 5 standard deviations above the mean^100^. We then removed candidate IEDs that a) had amplitudes below 3 standard deviations of the mean in the unfiltered LFP data, b) were longer than 200 ms, and c) occurred within 250 ms of another IED. To systematically identify non-IED artifacts (sharp, high-amplitude transients resulting from muscle artifact, cable movement, electronic interference, etc.), we a) computed the gradient of the unfiltered LFP, b) z-scored the resulting signal, and c) identified peaks in this signal that were at least 5 standard deviations above the mean. For the resulting IEDs and sharp transients, we removed the time-points that adhered to these criteria as well as 100 ms surrounding each artifactual event from the LFP data prior to computing power to mitigate their influence.

We synchronized the neural data to the behavioral task using a photodiode attached to the laptop screen that recorded screen updates in the behavioral task, recorded in the electrophysiological system as an analog input (Mt. Sinai) or a TTL pulse generated by the computer during the onset of specific events in the task (University of Iowa). After synchronization, we split the continuous LFP data into epochs surrounding behavioral events of interest, with a 1000 ms buffer used to limit edge effects in power and connectivity computations. We localized the electrodes by co-registering the preoperative magnetic resonance imaging (MRI) with the postoperative computed tomography (CT) scans using the Statistical Parametric Mapping (SPM) toolbox in MATLAB through a graphical user interface for locating intracranial electrodes (LeGUI)^101^. We used LeGUI to automatically identify electrodes in the co-registered imaging, and then manually corrected the electrode locations and identified electrodes that were outside of the brain or in white matter. We assigned each electrode a location using the Yale Brain Atlas^102^.

### Multivariate encoding of RPE

We computed time-frequency representations (TFR) for each event of interest, in each electrode, using Morlet wavelets^103^, with 30 log-spaced frequencies between 2-200 Hz, with 30 corresponding log-spaced cycles from 3-10 cycles. We baselined the resulting power values using the TFR computed from the fixation cross period, by computing the mean and standard deviation of power at each frequency during the fixation cross and using these values to z-score power during each event of interest.

We quantified the temporal evolution of RPE-related neural representations using multivariate pattern analysis of high-frequency activity (HFA). For each participant and anatomically defined region of interest (ROI), HFA was averaged across the 70–200 Hz frequency range, smoothed using a 200-ms sliding window, and downsampled by a factor of 5 before classification. Pattern classification was performed separately within each participant and ROI, using the pattern of HFA across electrodes at each time point as the input to the classifier.

For within-period decoding, we trained linear discriminant classifiers to distinguish among three RPE categories during the feedback period. Trial-wise model-derived RPE values were divided into terciles to define negative, neutral, and positive RPE categories, which served as class labels for multiclass linear discriminant analysis with shrinkage regularization. Classification performance was quantified as decoding accuracy using five-fold cross-validation at each time point. Statistical significance was assessed at the group level using permutation-based, cluster-corrected procedures, with trial labels shuffled within participants to generate null distributions. Contiguous time points exceeding a cluster-forming threshold of t = 1 were grouped into clusters, and cluster-level significance was assessed against the permutation distribution to control for multiple comparisons across time.

### Reactivation of multivariate RPE representations during memory

To test whether neural representations expressed during feedback re-emerged during subsequent memory retrieval, we performed cross-temporal, cross-period decoding between feedback-locked and memory-cue-locked activity. For each participant and ROI, classifiers were trained separately at each time point during the feedback period and tested at each time point during the subsequent memory-cue period, producing a two-dimensional generalization matrix in which each element represents decoding performance for a given feedback-training time and memory-testing time. This analysis tests whether distributed neural patterns associated with different RPE categories during feedback generalize to neural activity expressed during subsequent memory retrieval.

Statistical significance of the group-level generalization matrix was assessed using cluster-based permutation testing across the feedback-by-cue time matrix, using a cluster-forming threshold of t = 1. For significant clusters, each participant’s reactivation strength was quantified as the mean decoding accuracy within the significant cluster relative to chance performance. This subject-level measure was subsequently used for individual-differences analyses.

### Computing theta-phase synchrony

To quantify undirected and directed theta-band coupling between hippocampus (HPC) and anterior cingulate cortex (ACC), we estimated pairwise phase consistency (PPC) and the phase-slope index (PSI) on event-locked iEEG epochs. For each participant, anatomically labeled HPC and ACC contacts were pooled within subject.

PPC was estimated with multitaper spectral connectivity in the theta band (4-8 Hz), averaging across frequencies within the band to yield one PPC value per electrode pair^104^. Analyses were performed separately within reward-prediction-error (RPE) terciles (negative, neutral, positive). To ensure that connectivity estimates were not confounded by an electrode pair’s idiosyncratic theta prevalence, we generated 500 surrogate datasets for each condition by repeatedly cutting each channel’s epoch at a random time point and reversing the two temporal blocks, preserving univariate spectral structure while destroying genuine inter-regional dependence^105^. Observed PPC was then z-scored relative to the mean and standard deviation of each pair’s surrogate distribution.

To assess directed theta coupling, we computed trial-wise PSI between the same HPC (seed) and ACC (target) contacts using a continuous Morlet wavelet transform over 4-8 Hz (0.5 Hz steps; 4 cycles)^106^. Positive PSI indicates that HPC leads ACC. Estimates were then z-scored against 500 cut-and-reverse evoked surrogates generated as above, yielding a surrogate-normalized PSI-*z* for each trial and electrode pair.

### Identifying cross-regional reactivation during successful recall

We quantified cross-temporal representational similarity analysis (RSA) on trial-wise HFA aligned to encoding and retrieval. HFA signals were extracted from electrodes assigned to each ROI, smoothed with a 200-ms moving window, and downsampled by a factor of 5. For each subject, ROI pair, and memory condition (old vs. new) separately, we computed a representational dissimilarity matrix (RDM) at every encoding and retrieval time point by taking the Euclidean pairwise distance between multi-electrode trial-wise activity patterns. We then generated a cross-temporal RSA map by correlating the upper-triangular elements of the encoding and retrieval RDMs at every time-point combination using Spearman’s rho-a, yielding separate remembered and forgotten heat maps and a condition-difference map (remembered minus forgotten). To assess subject-level robustness, we constructed a permutation null by shuffling trial-wise memory labels 250 times, recomputing the difference heat map on each iteration, and estimating element-wise z-scores and two-sided empirical p values from the resulting null mean, null variance, and exceedance counts. For group inference, subject-level difference maps were stacked within each ROI pair and tested against zero using a cluster-based one-sample permutation procedure (5,000 permutations, one-tailed), and significant clusters were defined at cluster-level p≤ 0.05. We used a one-tailed test because we hypothesized that successful recognition produces greater representational similarity than unsuccessful recognition. This procedure thus isolates the extent to which trial-specific multivariate encoding patterns are reactivated during retrieval, and whether such reactivation is selectively stronger for remembered than forgotten items.

### Hippocampal ripple detection

We detected high-frequency ripple events on anatomically labeled HPC contacts. Putative interictal epileptiform discharges (IEDs) were identified first to exclude artifactual high-frequency bursts: for each electrode, the signal was bandpass-filtered at 25-60 Hz (zero-phase FIR), converted to an analytic envelope via the Hilbert transform, squared, and low-pass smoothed (40 Hz cutoff). The smoothed power was z-scored across all samples from that electrode, and contiguous excursions exceeding *z* > 4 were marked as IEDs at the sample of peak *z*. Ripple candidates were then detected on the same electrodes in the ripple band (70-180 Hz). After Hilbert envelope extraction, envelopes were robustly clipped at the median plus four median absolute deviations (scaled by 1.4826) to limit the influence of extreme bursts, squared, and smoothed as above. Events were initiated where smoothed power exceeded the mean plus four standard deviations (computed across all samples for that electrode), expanded until power fell below the mean plus two standard deviations, and merged when gaps were ≤30 ms. Candidate intervals shorter than 20 ms or longer than 200 ms were rejected. Event timing was assigned to the nearest trough of the 70-180 Hz bandpass-filtered voltage nearest the peak of smoothed power within the event. Finally, any candidate whose peak fell within 100 ms of a same-electrode, same-trial IED was excluded. Retained events were summarized as trial-wise ripple counts and rates (events per second of the analysis window) for downstream condition comparisons.

### Computing phase-amplitude coupling

We measured multivariate phase–amplitude coupling (PAC) between brain regions using a multivariate Gaussian-copula PAC approach. Coupling was computed separately for each trial and each theta–HFA frequency-bin pair. Rather than averaging pairwise electrode correlations, we estimated the mutual information (MI) between the multivariate phase vector (all phase-region channels in that hemisphere) and the multivariate amplitude vector (all amplitude-region channels in that hemisphere). Under the Gaussian copula model, MI was derived from the joint covariance of phase and amplitude variables across time points within the trial: separate covariance structures were computed for the phase set, the amplitude set, and the combined set. MI quantified the reduction in entropy of the amplitude vector given the phase vector. This yields a trial-specific comodulogram (5 phase bins × 5 amplitude bins) before trial-level averaging or condition comparisons.

PAC was computed separately for every trial. To assess whether coupling exceeded chance, we compared each trial’s PAC value to a null distribution built from 250 surrogate datasets generated by permuting the time structure of the amplitude signal. Each observed PAC value was converted to a standardized score by subtracting the mean of the surrogates and dividing by their standard deviation. For summary analyses, we averaged these standardized scores across the phase–amplitude frequency bins to obtain one trial-level coupling value per region pair and hemisphere.

### Statistical analysis and software

We conducted analyses in Python, using publicly available libraries. Electrode-level regression models were conducted using statsmodels, a Python library for statistical inference and linear modeling. Mixed-effects regressions were conducted using lmer in R.

## Acknowledgements

Research reported in this publication was supported by the internal funding from the Icahn School of Medicine at Mt. Sinai and the National Institutes of Health under award number K99/R00MH132873 to S.E.Q and award numbers R01MH122611, R01MH123069, R01MH124115, and R01DA060220 to X.G. The content is solely the responsibility of the authors and does not necessarily represent the official views of the National Institutes of Health. This work was supported in part through the computational and data resources and staff expertise provided by Scientific Computing and Data at the Icahn School of Medicine at Mount Sinai and supported by the Clinical and Translational Science Awards (CTSA) grant UL1TR004419 from the National Center for Advancing Translational Sciences. Research reported in this publication was also supported by the Office of Research Infrastructure of the National Institutes of Health under award number S10OD026880 and S10OD030463.

## Author Contributions

S.E.Q. and X.G. conceived the study; S.E.Q., F.P., L.N., A.R., H.K., C.K., C.G., B.D., and I.S. collected the data; S.E.Q. analyzed the data; and S.E.Q., I.S. and X.G. wrote the manuscript.

## Code Availability

Analysis code will be available upon publication at https://github.com/seqasim/MemoryBandit_EMU.

---

**Figure S1:**
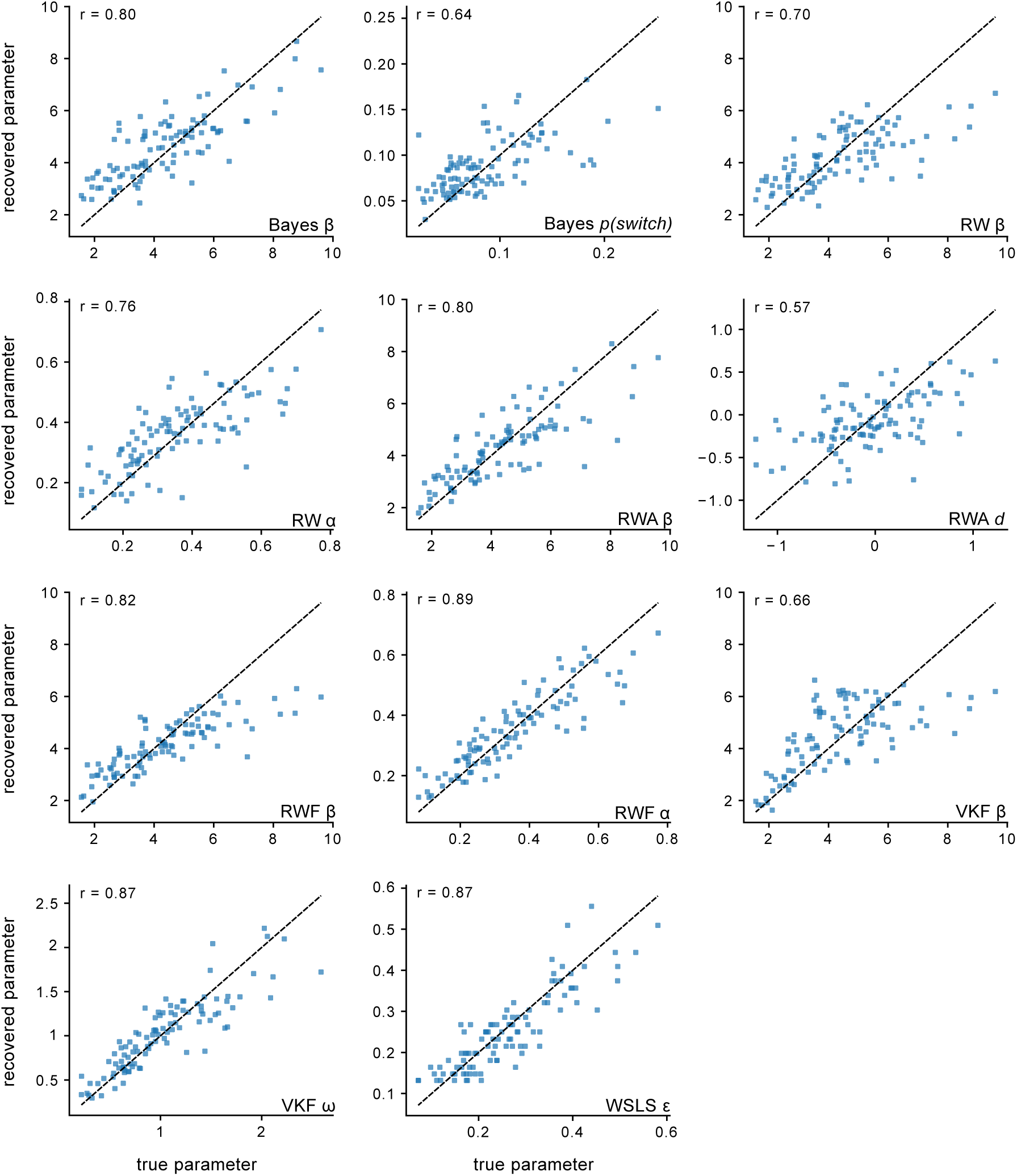
Parameter recovery for candidate reinforcement-learning models. Recovered vs. true parameter values from equal-design parameter recovery (*N* = 100 simulated subjects × 80 trials per model, matching the empirical task design), re-estimated via hierarchical expectation-maximization using the same production settings applied to real data. Each panel corresponds to one free parameter of one candidate model: Bayes (fixed-reward) inverse temperature (β) and switch probability (*p*(switch)); Rescorla–Wagner (RW) learning rate (α) and inverse temperature (β); valence-asymmetric RW (RWA) inverse temperature (β) and asymmetry (*d*); fictive RW (RWF) learning rate (α) and inverse temperature (β); volatility Kalman filter (VKF) inverse temperature (β) and volatility-learning rate (ω); and win-stay-lose-shift (WSLS) lapse rate (ε). Dashed lines indicate the identity line (perfect recovery); Pearson’s *r*between true and recovered values is reported in the top-left of each panel.

**Figure S2:**
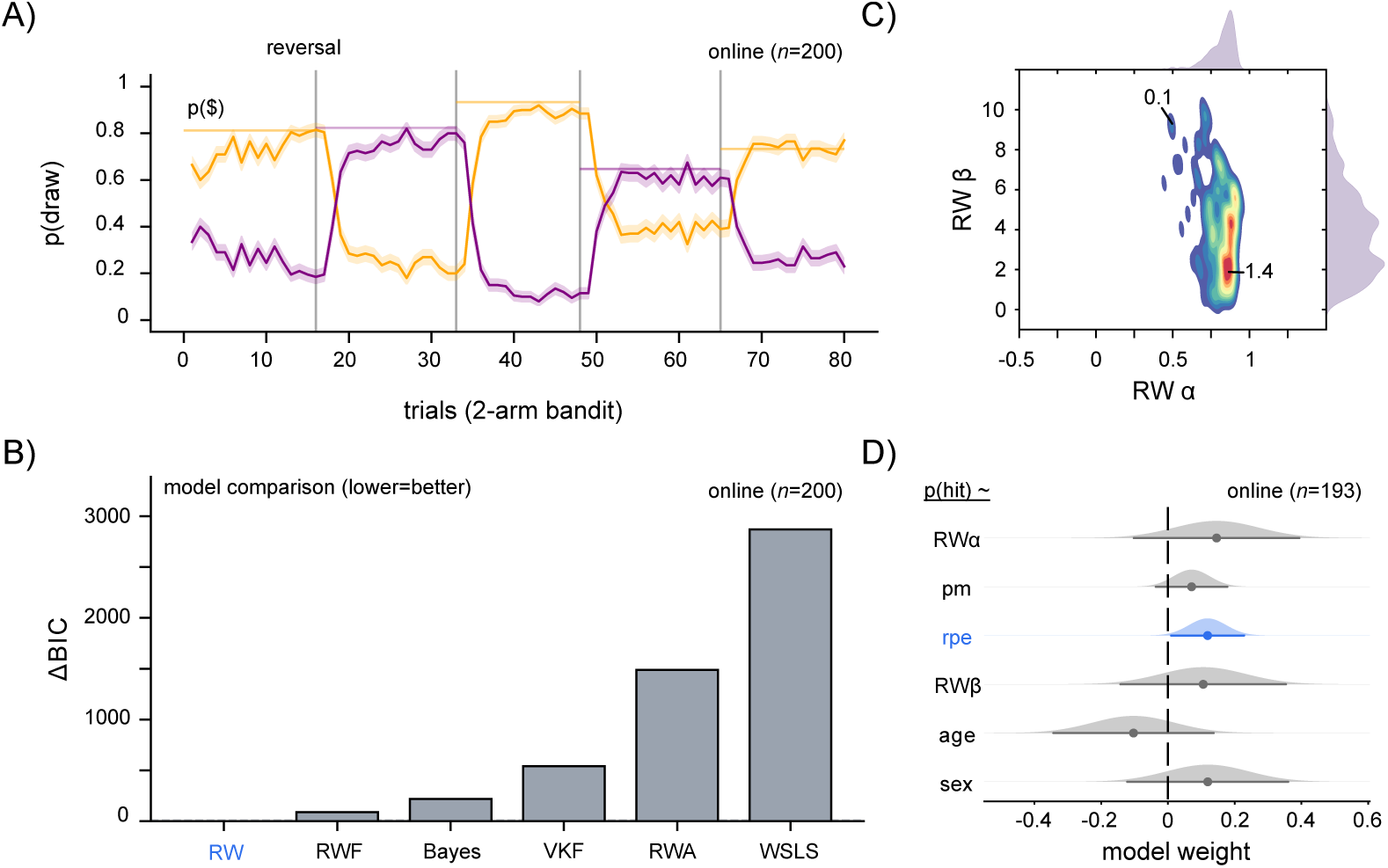
Healthy prolific cohort behavioral validation. A) Probability of drawing from the male (orange) or female (purple) decks as a function of trial and reward block (mean reversal trial indicated by vertical lines). Horizontal lines denote probability of reward for the more rewarding deck on each block. Shaded lines denote a 95% confidence interval. B) Model performance, specified by the integrated BIC across six candidate reinforcement-learning models fit hierarchically to subjects’ choice sequences on the 2-arm bandit task. Bar height represents ΔBIC, each model’s integrated BIC relative to the best-fitting model in the set (best = 0); lower is better. C) Joint and marginal distributions of parameter estimates from the RW model, depicting the learning rate (α) and inverse temperature (β). Warm colors indicate higher density. D) Sampling distributions for the fixed effects in a binomial mixed-effects model examining how trial-level reward prediction error (rpe), image memorability (pm), subject-level Rescorla–Wagner parameters (learning rate α, inverse temperature β), age, and sex influence recognition memory success in the online cohort (n=193). Each curve is a normal approximation to the coefficient’s sampling distribution, and the horizontal bar spans the 95% Wald confidence interval. The dashed vertical line indicates a coefficient of 0. Distributions whose 95% CI excludes 0 are shaded blue, indicating a statistically reliable effect, while those whose CI includes 0 are shaded gray.

**Figure S3:**
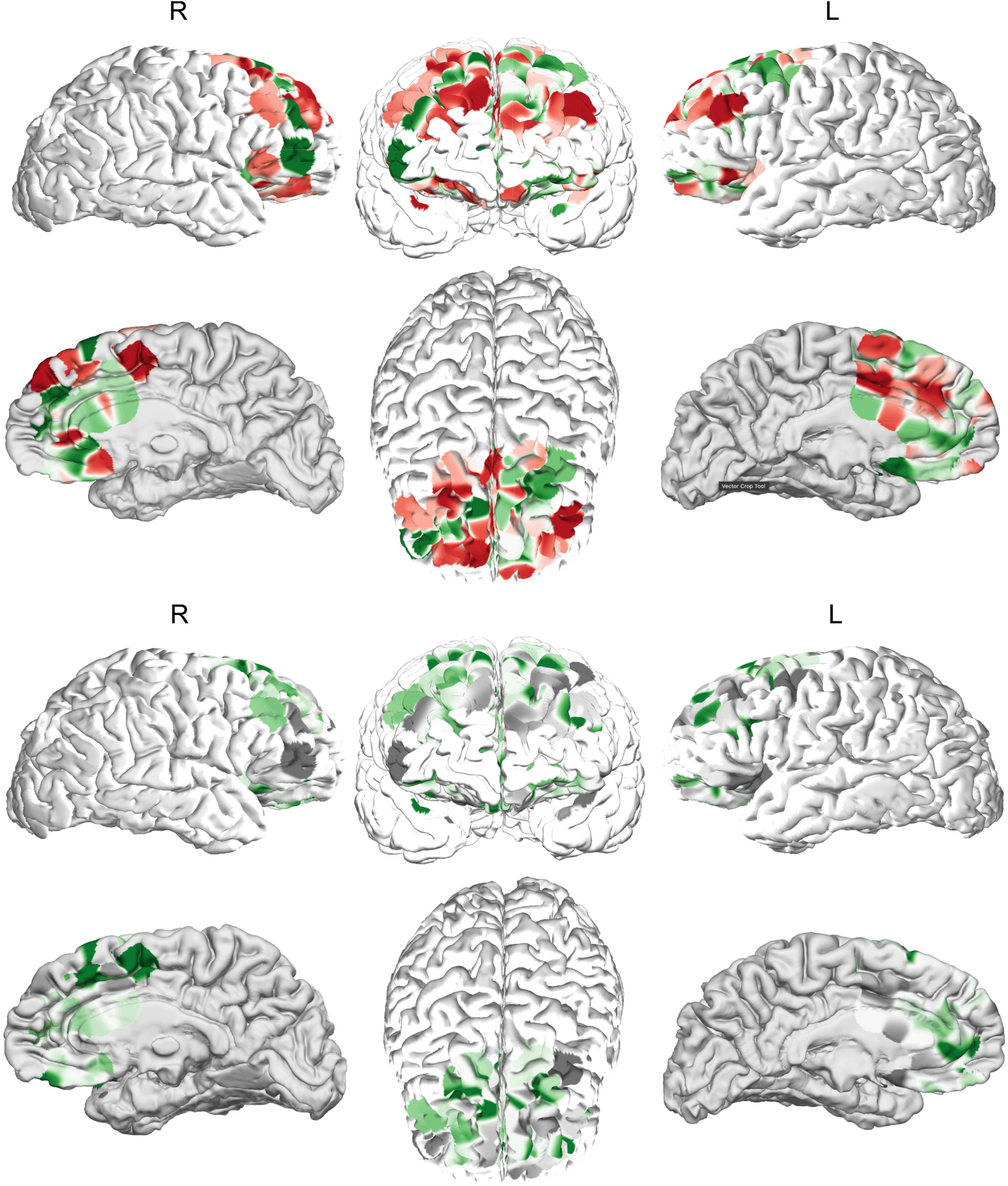
Prefrontal activation patterns for positive, negative, and neutral RPE encoding during feedback. Prefrontal activation patterns computed by multiplying the model weight vector by the feature covariance matrix, yielding a pattern that reflects how each electrode covaries with positive RPE compared to negative RPE (top) and neutral RPE (bottom). Timepoints for visualization were selected from the significant clusters for each individual ROI: OFC, ACC, dmPFC. Green denotes increased activation for positive RPEs, red denotes increased activation for negative RPEs (top), and gray denotes increased activation for neutral RPEs (bottom).

**Figure S4:**
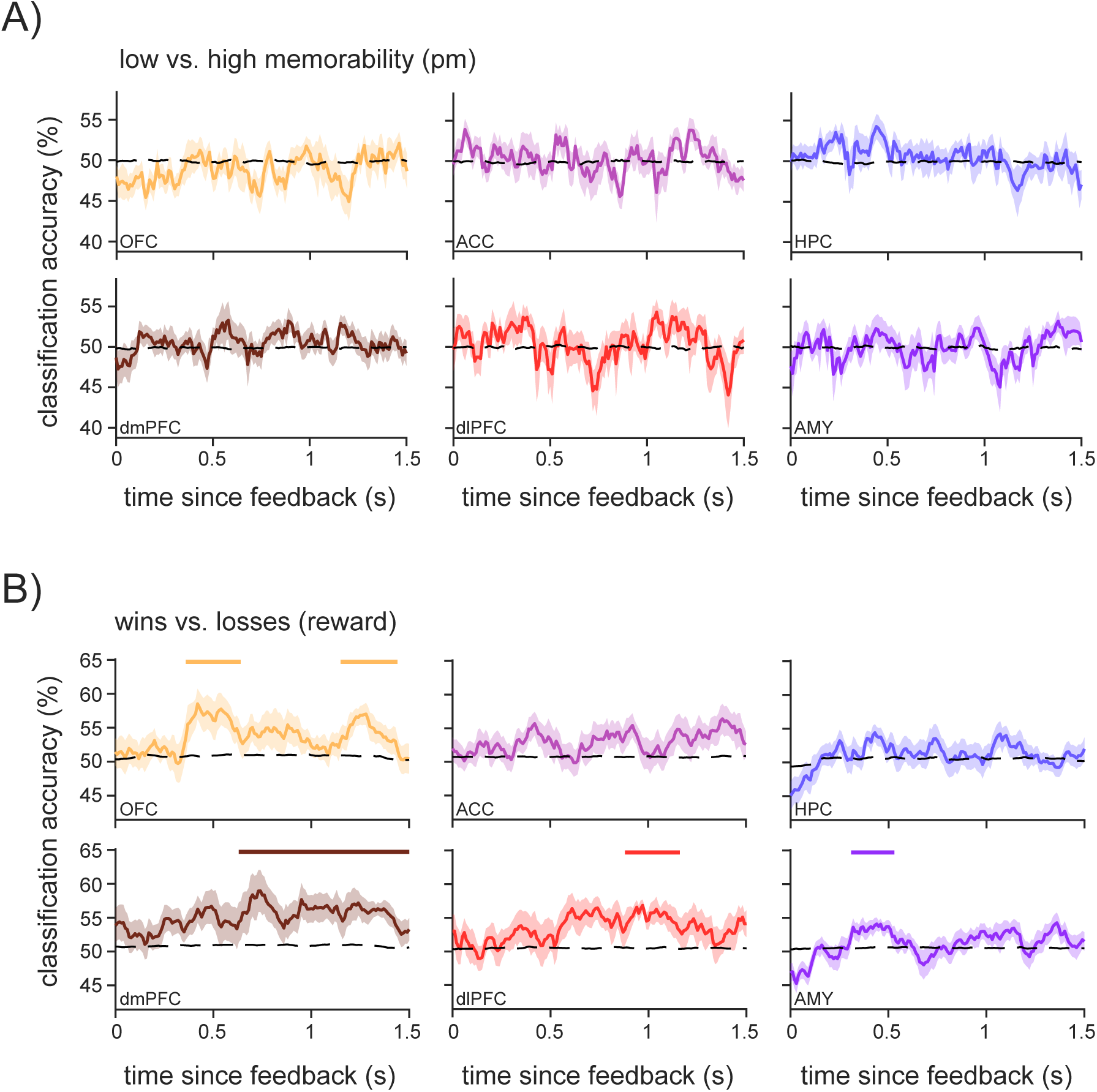
Perceptual memorability cannot be decoded during feedback. A) Cross-validated binary LDA classifier accuracy for memorability (high vs. low, median split) for each region of interest. The dotted line denotes the mean of the surrogate values. Solid lines denote significant time clusters computed using shuffled null data. Shaded lines indicate standard error across subjects. B) Cross-validated binary LDA classifier accuracy for reward outcome (wins vs. losses) for each region of interest. The dotted line denotes the mean of the surrogate values. Solid lines denote significant time clusters computed using shuffled null data. Shaded lines indicate standard error across subjects.

**Figure S5:**
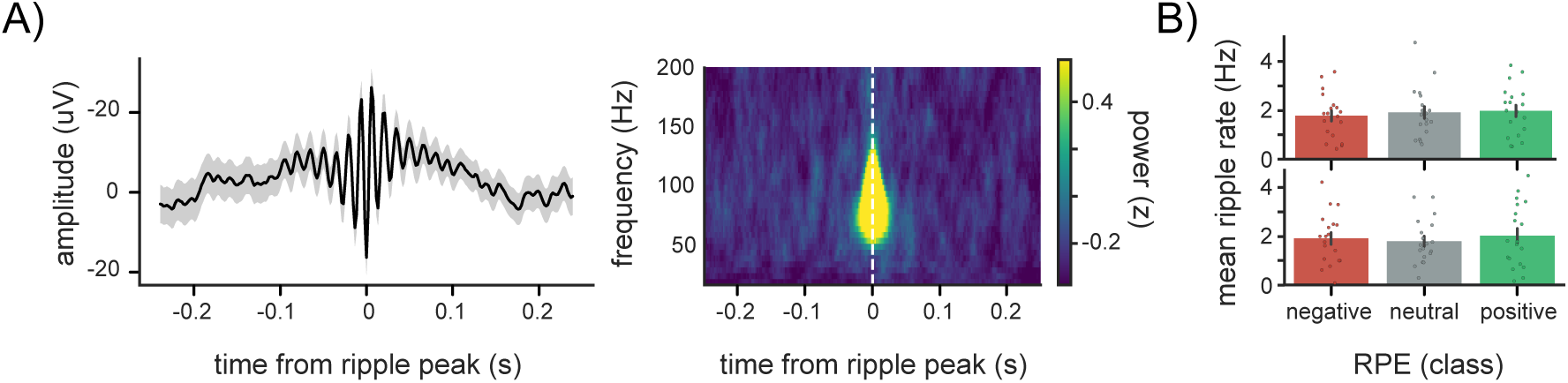
**Hippocampal ripple rate does not di**ff**er as a function of RPE.** A) Left: example raw LFP trace of grand-average ripple (one subject; 340 ripples) in the hippocampus. Solid line denotes the mean across ripples. Shaded line denotes standard error. Right: spectrogram of the detected ripples in the same example subject. Yellow denotes higher power. B) Comparison of ripple rates between feedback (top) and recognition cue periods (bottom). Bar heights denote the mean hippocampal ripple rate across subjects (dots). Error bars denote standard error.

**Figure S6:**
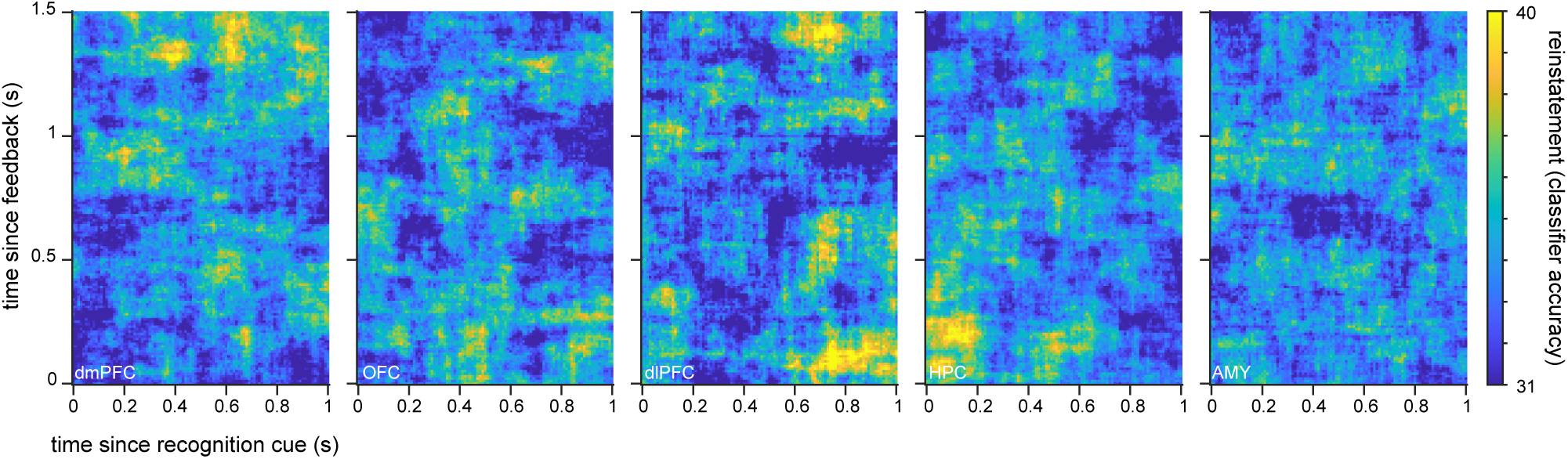
Cross-decoding for other ROI. Performance of cross-decoding classifier for RPE class trained on the feedback period (y-axis) and tested on the recognition cue period (x-axis), for all other regions of interest. Warmer colors indicate higher classifier performance.

**Figure S7:**
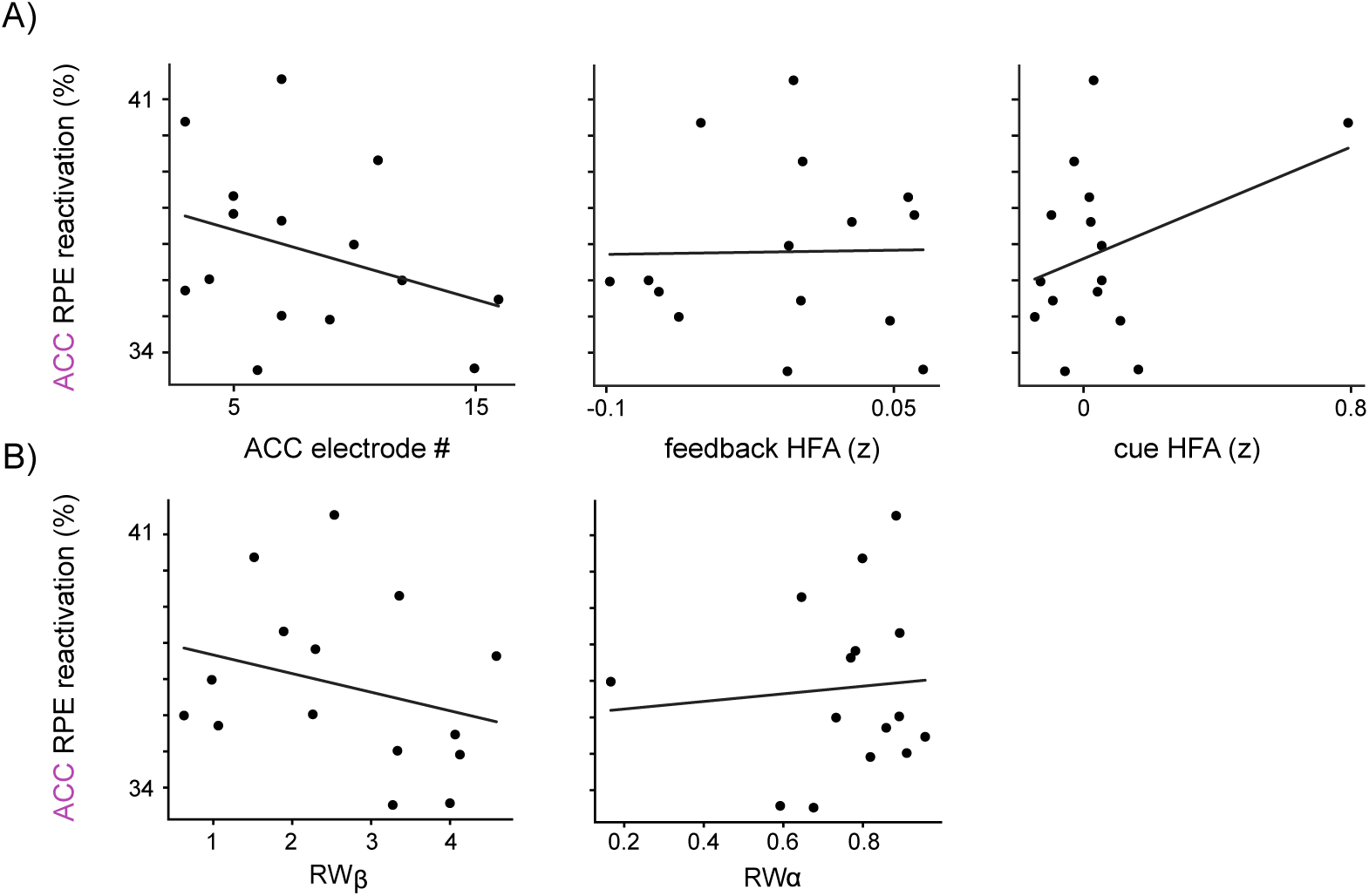
ACC reinstatement not explained by subject-level covariates. A) Scatter plots of the relationship between ACC reactivation in the significant cluster and: the number of ACC electrodes (left), overall ACC HFA during the feedback period (middle), and overall ACC HFA during the cue period (right). B) Scatter plots of the relationship between ACC reactivation in the significant cluster and: the learning rate (left), and inverse temperature (right).

**Figure S8:**
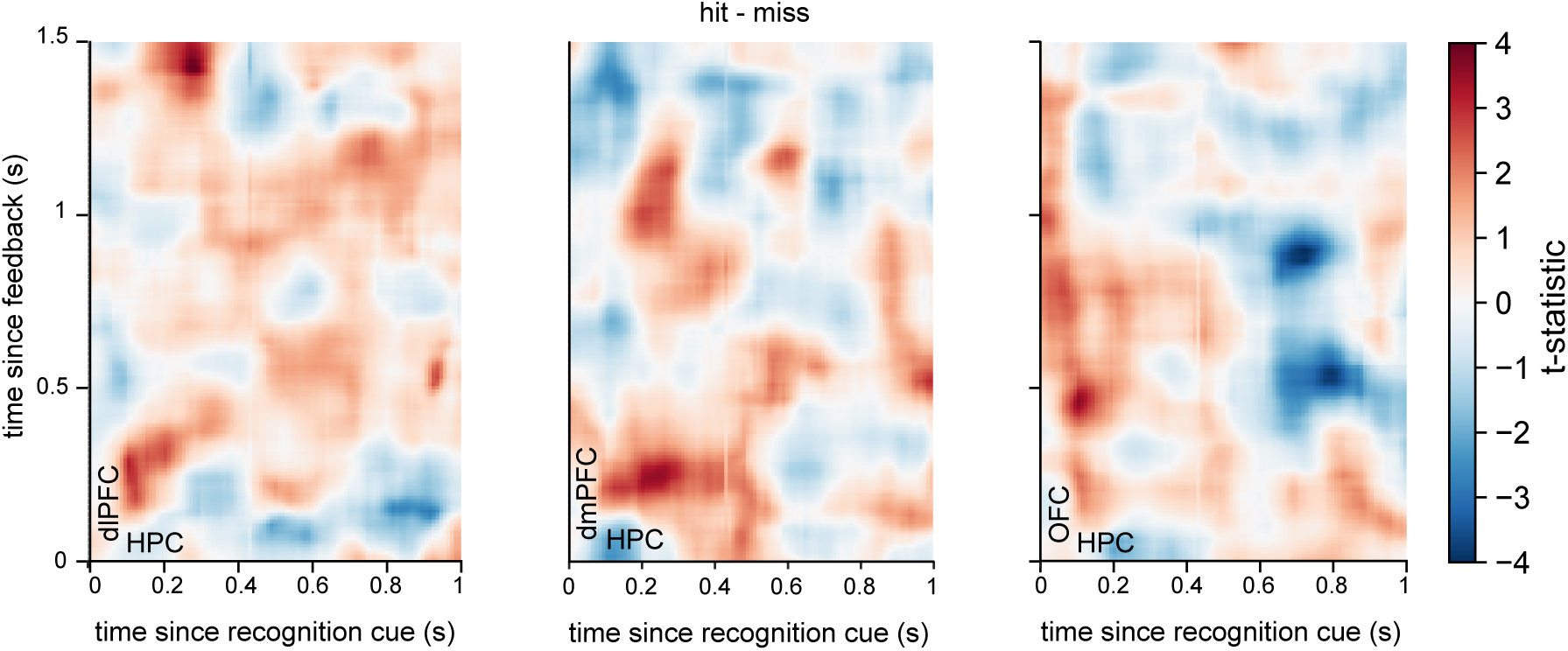
Between-ROI RSA analyses. Contrast heatmap of z-scored representational similarity (RSA) for all feedback and cue timepoints (hits vs. misses) for between-ROI analysis for dlPFC, dmPFC, and OFC feedback activity compared to HPC cue activity. Warmer colors denote higher RSA for hit trials compared to miss trials. Outline denotes significant cluster using permutation-cluster test.

**Figure S9:**
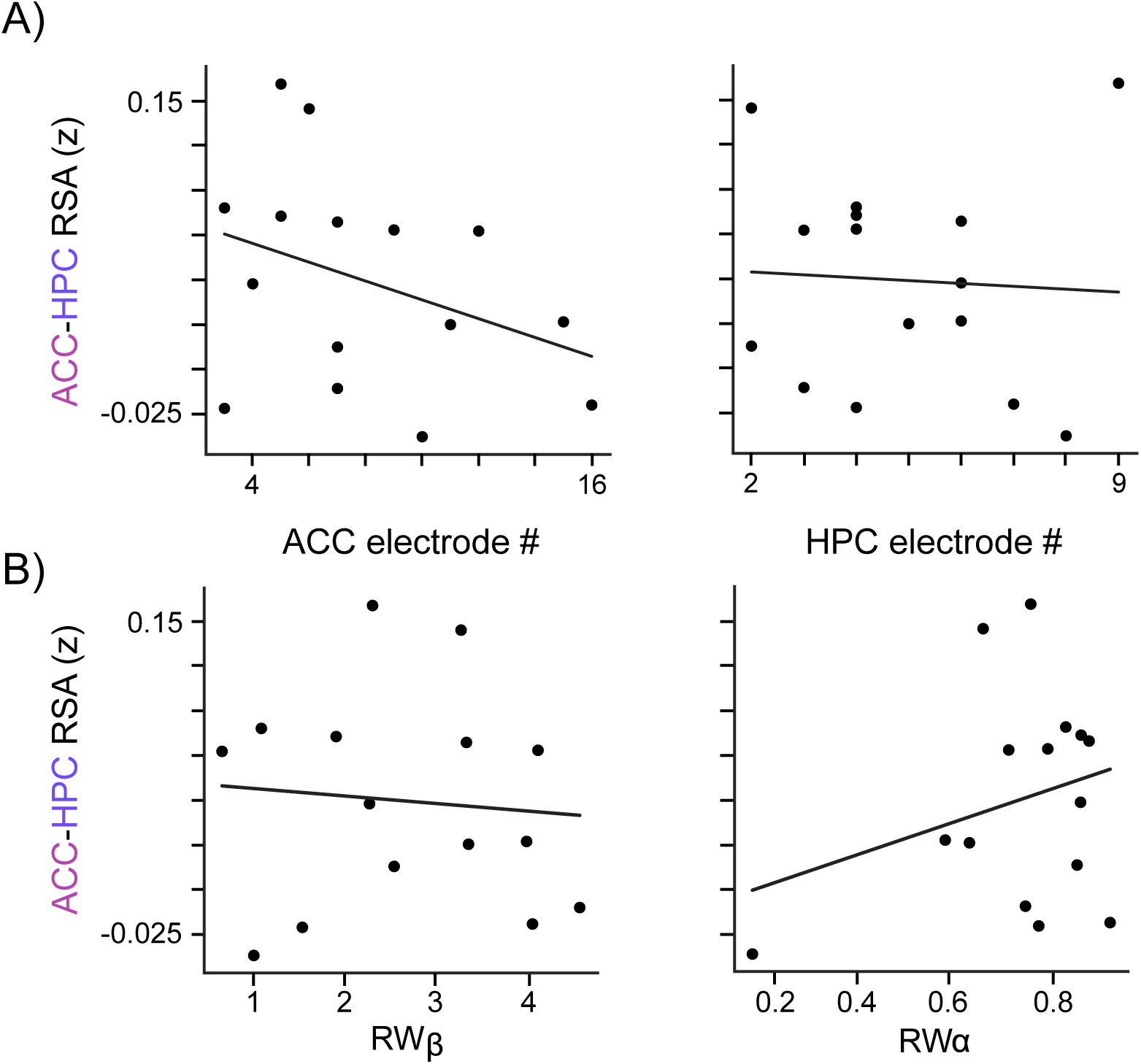
HPC reinstatement of ACC feedback activity not explained by subject-level covariates. A) Scatter plots of the relationship between ACC-HPC RSA and: the number of ACC electrodes (left), the number of HPC electrodes (right). B) Scatter plots of the relationship between ACC-HPC RSA and: the learning rate (left), and inverse temperature (right).

**Figure S10:**
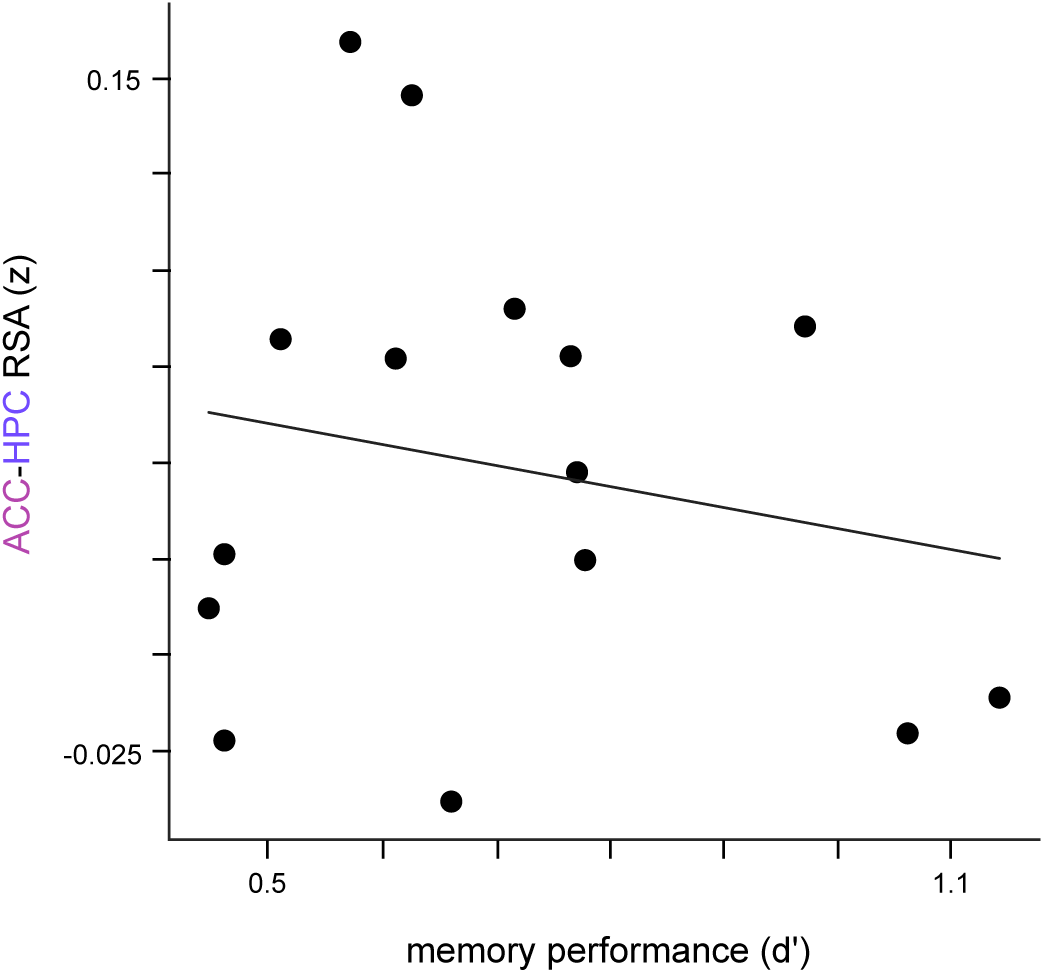
HPC reinstatement of ACC feedback activity not explained by memory performance. Scatter plots of the relationship between ACC-HPC RSA and memory performance.

**Figure S11:**
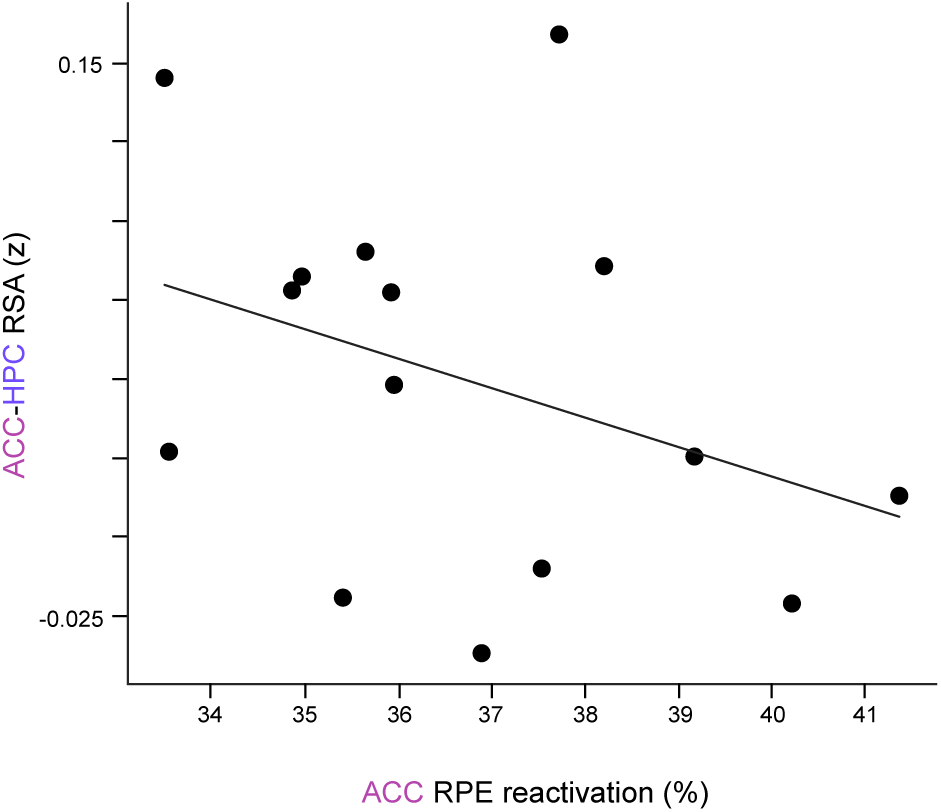
HPC reinstatement of ACC feedback activity not explained by ACC RPE reactivation. Scatter plot of the relationship between ACC reactivation in the significant cluster and ACC-HPC RSA in the significant cluster.

**Figure S12:**
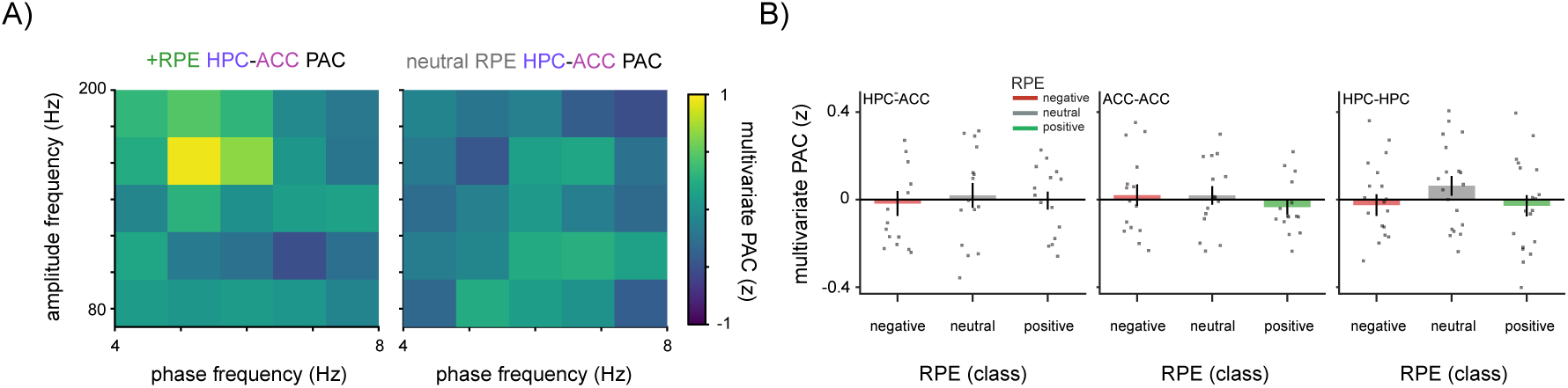
No evidence that RPE induces coupling between theta phase and HFA amplitude in or between regions. A) Example of multivariate phase-amplitude coupling computed between hippocampal theta phase and ACC HFA amplitude during the recognition cue period for positive RPE images (left) and neutral RPE images (right). Brighter colors denote larger z-scored phase-amplitude coupling. B) Mean phase-amplitude coupling during the cue period for negative, neutral, and positive RPE images for HPC-ACC (left), ACC-ACC (middle), and HPC-HPC (right). Bar height denotes mean across subjects (dots).

**Figure S13:**
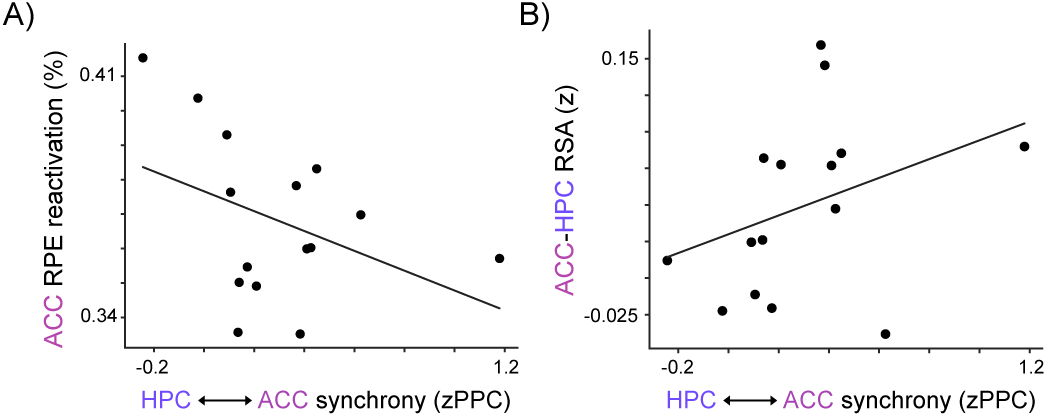
ACC reinstatement and HPC reinstatement of ACC feedback activity not explained by HPC-ACC synchrony. B) Scatter plot of the relationship between HPC-ACC theta synchrony during positive RPE cue trials and ACC reinstatement in the significant cluster. B) Scatter plot of the relationship between HPC-ACC theta synchrony during positive RPE cue trials and ACC-HPC RSA in the significant cluster.

